# Multipartite complexity of the lichen symbiosis revealed by metagenome and transcriptome analysis of *Xanthoria parietina*

**DOI:** 10.1101/2024.08.16.608140

**Authors:** Gulnara Tagirdzhanova, Klara Scharnagl, Neha Sahu, Xia Yan, Angus Bucknell, Adam R. Bentham, Clara Jégousse, Sandra Lorena Ament-Velásquez, Ioana Onuț-Brännström, Hanna Johannesson, Dan MacLean, Nicholas J. Talbot

## Abstract

Lichens are composite symbiotic associations of fungi, algae, and bacteria that result in large, anatomically complex organisms adapted to many of the world’s most challenging environments. How such intricate, self-replicating lichen architectures develop from simple microbial components remains unknown because of their recalcitrance to experimental manipulation. Here we report a metagenomic and metatranscriptomic analysis of the lichen *Xanthoria parietina* at different developmental stages. We identified 168 genomes of symbionts and lichen-associated microbes within a lichen thallus, including representatives of green algae, three different classes of fungi, and 14 bacterial phyla. By analyzing occurrence of individual species across lichen thalli from diverse environments, we defined both substrate-specific and core microbial components of the lichen. Meta-transcriptomic analysis of the principal fungal symbiont from three different developmental stages of a lichen, compared to axenically grown fungus, revealed differential gene expression profiles indicative of lichen-specific transporter functions, specific cell signalling, transcriptional regulation and secondary metabolic capacity. Putative immunity-related proteins and lichen-specific structurally conserved secreted proteins resembling fungal pathogen effectors were also identified, consistent with a role for immunity modulation in lichen morphogenesis.

## Introduction

Symbiosis is one of the most widespread and successful lifestyle strategies for biological organisms. It was first discovered in the form of lichens: long thought to be a single organism, lichens were revealed instead to be a stable relationship between a fungus and one or multiple photosynthetic microorganisms^1^. A unique and defining feature of the lichen symbiosis is a new body plan that arises only from the interaction. Stable and self-replicating over generations, the lichen phenotype does not resemble that of any of the symbionts grown in isolation. Lichen symbionts interact to create a single body (a thallus), which is often structurally complex and organized into multiple tissue-like layers. A major role in lichen development is believed to belong to the mycobiont – the fungal symbiont which contributes the vast majority of lichen biomass. Interwoven and glued together with extracellular matrix, mycobiont hyphae create tough outer layers of the lichen thallus, with photosynthetic symbionts (photobionts) typically inhabiting the layer beneath, where they can take advantage of sunlight^2^. In addition to the mycobiont and photobiont, many lichens contain additional microorganisms, chiefly bacteria and yeasts, at least some of which are stably associated with lichens^3,4^.

The molecular mechanisms required for lichen development and growth remain unknown. While we can hypothesize that some may be similar to those involved in development of complex fungal structures, such as mushrooms, this hypothesis needs to be tested, and we also need to explain the remarkable coordination of growth between symbionts. The reason behind such limited knowledge of lichen symbiotic development lies in the recalcitrance of lichens towards laboratory experimentation. Individual symbionts often grow extremely slowly^5^ and, with one exception, have never been genetically modified. The only exception, the mycobiont of *Umbilicaria* lichens, is highly unusual in its dimorphic growth habit^6^, which makes it easier to manipulate, but also raises questions of whether its study is applicable to other lichens. Lichen phenotypes cannot be recreated from axenic cultures in the lab, leaving us with no mechanistic insight into lichen development.

In this report, we use metagenomics and metatranscriptomics to characterize a lichen symbiosis and identify processes involved in symbiosis maintenance and development. As a model, we used *Xanthoria parietina* – a widespread lichen that has served as a model system in studies of lichen anatomy and population genetics^7,8^. *X. parietina* is believed to have no vertical co-transmission of symbionts, which disperse on their own. Hence, germinating sexual spores of the mycobiont must establish connection to a *Trebouxia* photobiont and, potentially, other members of the lichen microbiota every time a new lichen forms. We establish *X. parietina* as our model system by analyzing the genome of its mycobiont and by characterizing the diversity of microorganisms present in lichen samples. We compare mycobiont gene expression between intact lichen thalli and lab cultures, and, for the first time, compare different developmental stages of a lichen thallus in order to identify genes and molecular processes involved in lichen morphogenesis. Finally, we perform the first in-depth analysis of a lichen mycobiont secretome and identify potential symbiosis-associated lichen effector proteins.

## Results

### Organisation of the *Xanthoria* mycobiont genome

We first generated a reference genome of the *X. parietina* mycobiont. Long-read metagenomic data from a *X. parietina* thallus collected at the Norwich Research Park (Figure 1A) yielded a high-quality genome assembly of the mycobiont. Data were assembled and binned to remove sequences from any organism other than the mycobiont. The final mycobiont genome assembly consisted of 58 contigs for a total of 29.96 Mbp; the N50 of the assembly equaled 1.59 Mbp (Figure 1B). The assembly had completeness scores of 96.1% according to BUSCO5 and of 98.1% according to EukCC2, where completeness score is defined as 100% minus the percentage of missing markers (Figure 1C). *De novo* annotation of the genome resulted in 10,727 gene models and 11,185 transcripts. The genome size and completeness and annotation statistics are consistent with other high-quality genomes from the class Lecanoromycetes published to date^9–12^. In the genome, we identified 59 biosynthetic gene clusters (Data S1).

The repeat content of the genome was 12.7%, with long terminal repeat elements accounting for nearly half of the repeated component of the genome (Data S1). Repeats were not evenly spread across contigs and instead formed regional clusters, which corresponded to genome regions with lower GC content (Figure 1B). By screening the genome for signatures of Repeat Induced Point Mutation (RIP)^13^, we discovered that these low-GC/high-repeat regions can also be considered Large RIP Affected Regions (LRARs) (Figure 1B). In total, we identified 158 LRARs that account for 8.5% of the genome (Data S1).

To predict the ploidy level of the *Xanthoria* mycobiont, we analyzed minor allele frequency (MAF) distribution in eight metagenomic samples (see below) using the newly produced mycobiont genome for variant calling. While some samples showed haploid-like patterns, other patterns had unusually high numbers of peaks in the distribution (Figure S1), consistent with neither haploid, diploid, or triploid signals.

**Figure 1.**
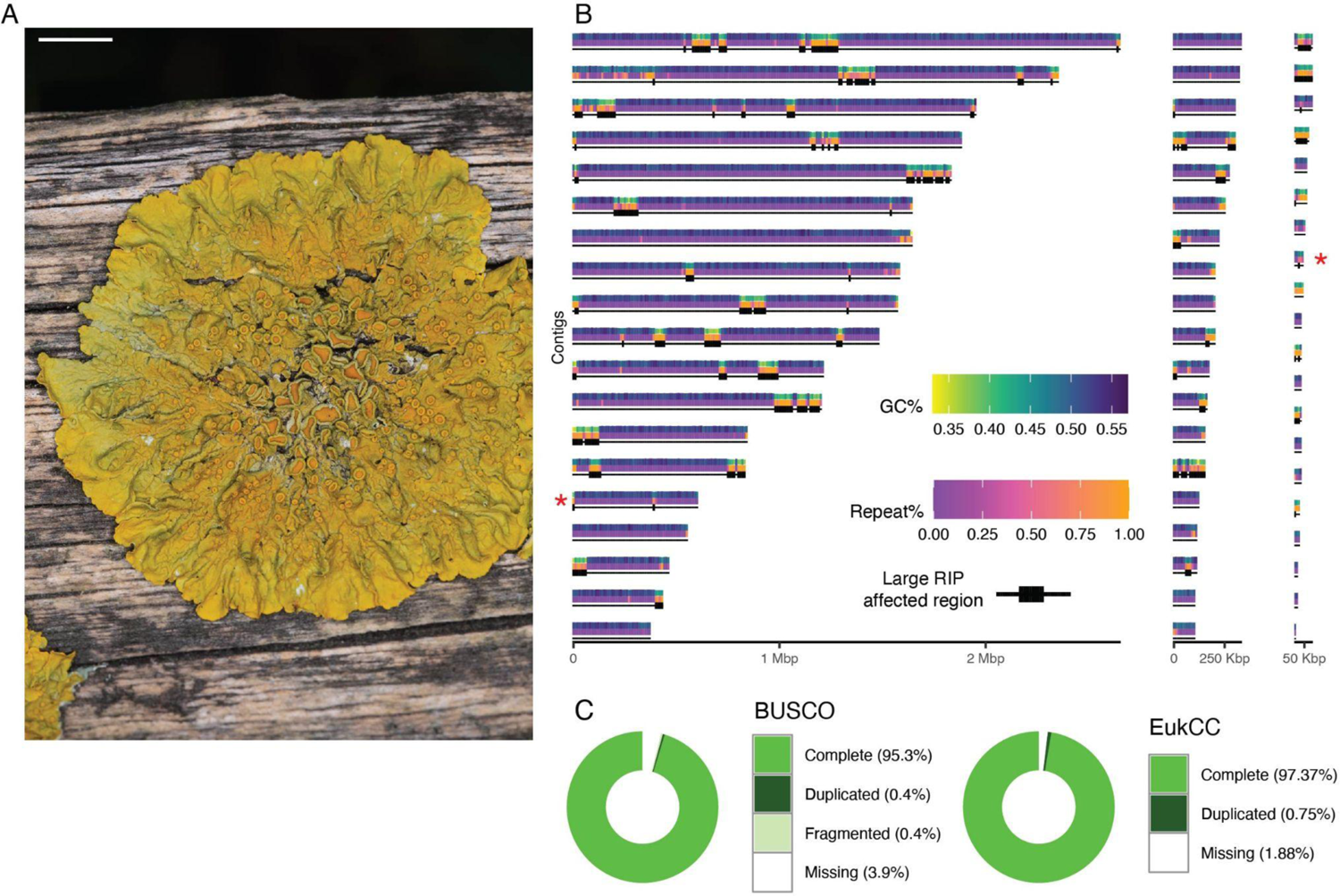
The genome of *Xanthoria parietina* mycobiont. A. *X. parietina* thallus on wood. Scale bar 5 mm. B. *X. parietina* mycobiont nuclear genome. Each contig is represented by three annotation tracks: GC content, repeat content, and presence of Large RIP Affected Regions (LRARs). The x-axis corresponds to contig length. Red asterisks show telomeric repeats. C. Genome completeness scores estimated by BUSCO5 (ascomycota_odb10 database) and EukCC2 (NCBI 78060 Parmeliaceae database). See also Figure S1 and Data S1.

### 168 metagenome-assembled genomes (MAGs) can be isolated from *Xanthoria* metagenomes

To characterize the organismal composition of *X. parietina* thalli and account for all species detected in shotgun sequencing experiments, we generated eight deeply-sequenced metagenomes (minimum of 24.7 Gbp of raw data and 85x mycobiont genome coverage) from samples of *X. parietina* collected from Norwich Research Park from different substrates and growth conditions: concrete (n=3), tree bark (collected fresh, n=3), and tree bark (incubated in a growth chamber for 18 months, n=3). From these metagenomes, we extracted and annotated 168 medium and high-quality non-redundant MAGs each corresponding to a distinct species-level lineage (Data S2; see Methods for details of MAG filtering and dereplication). All eleven eukaryotic MAGs belong to either fungi or algae. The seven fungal MAGs include the *X. parietina* mycobiont (Figure 2A, Figure S1) and three distantly related mycobionts of other lichen symbioses, from classes Lecanoromycetes and Lichinomycetes. These genomes were likely obtained due to propagules of these fungi on the surface of *X. parietina* samples. In addition, three MAGs of Chaetothyriales (Eurotiomycetes), a group of black yeasts reported from various lichens as potential endophytes or parasites^14–16^, were detected in three of the eight *Xanthoria* samples. All four algal MAGs belonged to different strains of *Trebouxia* (Figure 2B), the previously reported photobiont of *X. parietina* lichen^7^. The remaining 157 MAGs are shared between 14 bacterial phyla, with 59% from just two phyla: Proteobacteria and Actinobacteriota (Data S2, Figure 2C). The two bacterial genera with most MAGs were *Sphingomonas* (Sphingomonadaceae, Alphaproteobacteria; n=18) and clade CAHJXG01 (Acetobacteriaceae, Alphaproteobacteria; n = 9) (Data S2).

**Figure 2.**
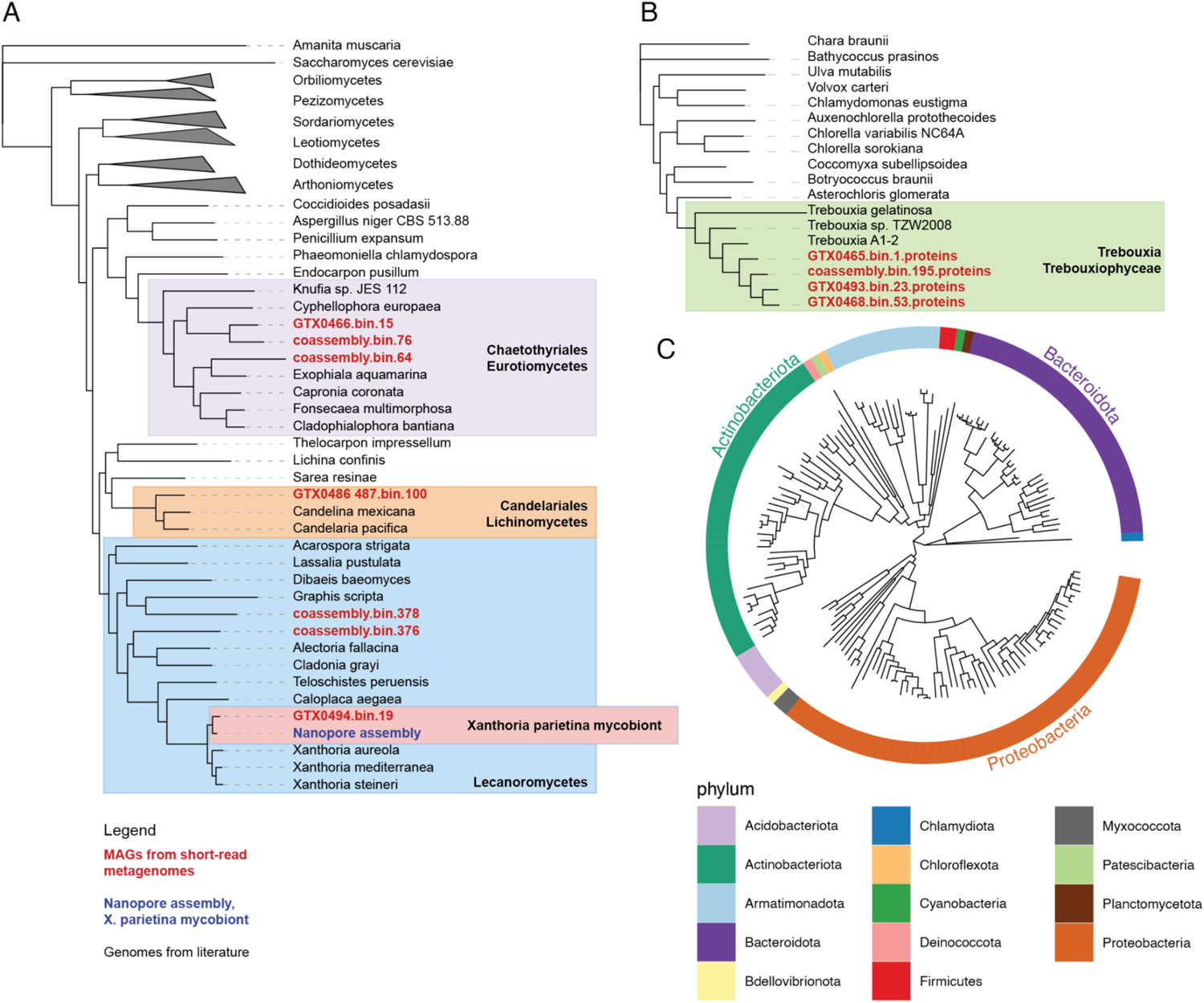
Maximum-Likelihood trees of genomes assembled from *X. parietina* metagenomes. A. Fungal phylogenomic tree. Metagenome-assembled genomes (MAGs) assembled from the eight metagenomes shown in red, long-read genome assembly shown in blue. To clarify the taxonomic position of MAGs, we added reference genomes with known identities (shown in black; Data S2). The genome of *Amanita muscaria* was used as an outgroup to root the tree. B. Algal phylogenomic tree. MAGs assembled from metagenomes shown in red, reference genomes shown in black (Data S2). The genome of *Chara braunii* was used as an outgroup. C. Bacterial phylogenomic tree. The color track shows taxonomic assignment.

Next, we mapped the presence/absence of each MAG across eight *X. parietina* samples by mapping metagenomic reads onto the MAG catalog. We compared lichen samples collected from different substrates: concrete, tree bark (collected fresh), and tree bark (incubated in a growth chamber for 18 months). Clustering lichen samples based on the occurrence matrix revealed that samples collected from lichens growing on concrete differed from bark samples (Figure 3A). Concrete samples also had the highest number of unique MAGs (Figure 3B). The role of growth substrate in determining taxonomic composition of lichen-associated microorganisms is also confirmed by our analysis of an additional sample of a different *Xanthoria* species, *X. calcicola*, collected from concrete. This sample clustered with *X. parietina* samples from concrete and shared the majority of lineages present in these samples (Figure S2). At the same time, differences between substrates slightly decreased when diversity was considered at higher taxonomic levels: the percentage of lineages present in all three substrate types, for instance, was 19% for species-level lineages and 30% for family-level lineages (Figure 3B).

In addition to lineages present occasionally or in one type of substrate only, we detected generalist lineages present in all surveyed lichen thalli. As expected, a mycobiont MAG was detected in all samples and at least one MAG was assigned to the *Trebouxia* photobiont. The four detected photobiont lineages often co-occurred in various constellations; photobiont identity did not appear to depend on substrate (Figure 3C). We also detected 13 bacterial MAGs universally present, of which four came from *Sphingomonas* (Figure 3D, Data S2). Each metagenomic sample included at least six different *Sphingomonas* MAGs which were not substrate-dependent (Figure 3D). By contrast, CAHJXG01 showed substrate-dependency. While every sample contained at least one CAHJXG01 MAG, none of the MAGs were present universally. Instead, they formed two clusters based on substrate (Figure 3E). Both *Sphingomonas* and CAHJXG01 are frequent in lichens^17^, and other generalist bacteria have been reported from lichens too^18–20^. We conclude that lichen thalli contain a large number of associated microorganisms, that can be putatively split into a substrate-dependent lichen microbial community and a core lichen community.

**Figure 3.**
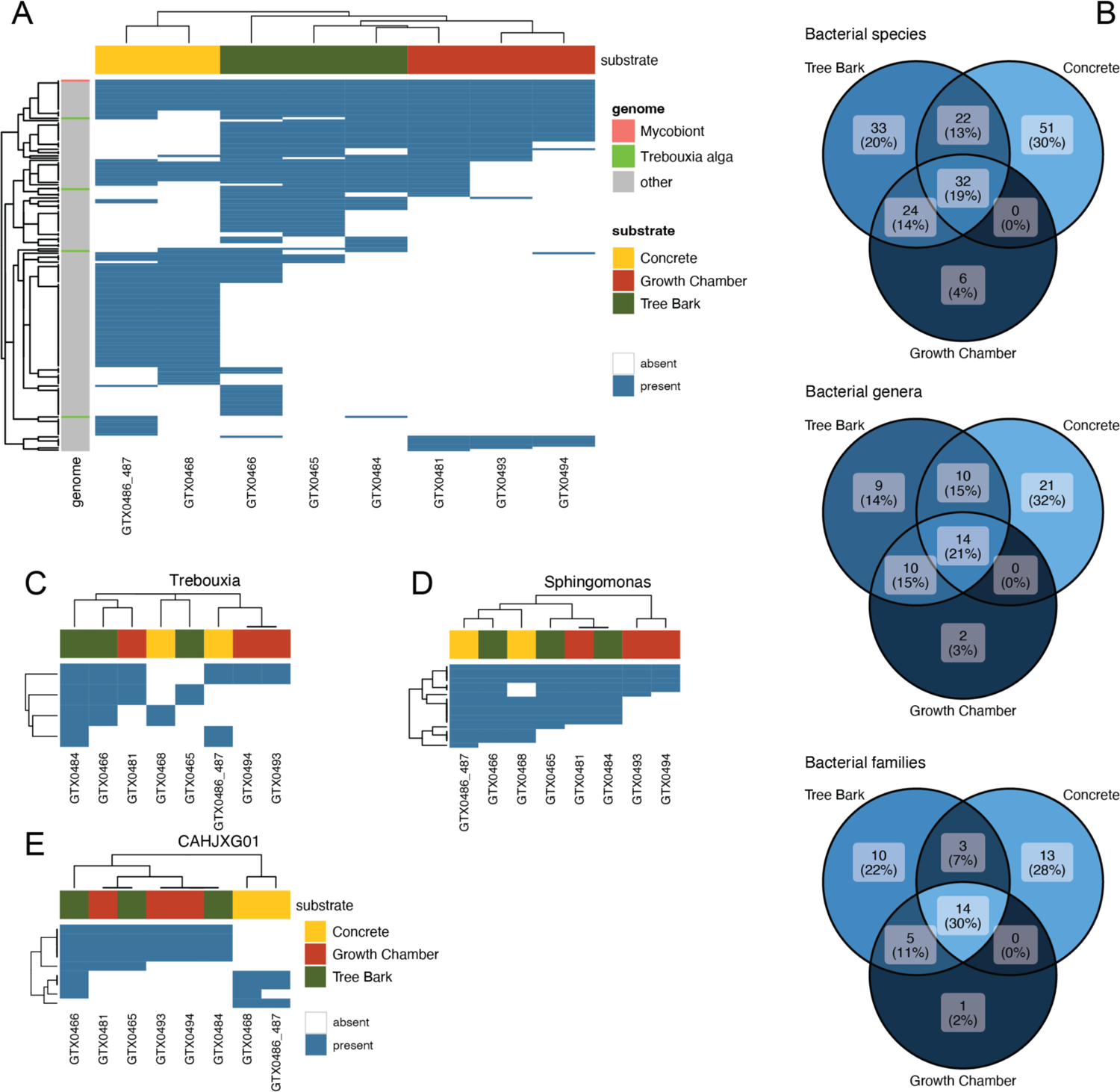
Species diversity detected in *X. parietina* metagenomes. A. Presence/absence map of 168 metagenome-assembled genomes (MAGs) assembled from *X. parietina* in eight metagenomes, divided by substrate. B. Venn diagrams showing shared and unique bacterial taxa (on the level of species = MAG, genus, and family) detected in *X. parietina* metagenomes from different substrates. C-E. Presence/absence map of selected lineages: C. Algae; D. *Sphingomonas*; E. Acetobacteraceae clade CAHJXG01. See also Figure S2.

### Differentially expressed genes disproportionally lack functional annotation and come from lichen-enriched orthogroups

To identify cellular processes involved in the lichen symbiosis, we compared gene expression of the mycobiont between intact lichen thalli at distinct developmental stages (17 samples from seven thalli; see below) and axenically grown mycobiont in pure culture (12 samples from four timepoints; Figure 4A). We pseudoaligned RNA-seq data to the reference, produced by compiling predicted transcriptomes from the mycobiont and non-mycobiont MAGs isolated from *X. parietina* metagenomes. The majority of reads in all libraries were aligned to the mycobiont transcriptome (Figure 4B), and because the mycobiont is responsible for >90% of the biomass of the lichen thallus^21^, we focused on its gene expression. Analysis of gene expression in the photobiont proved impossible at this stage, due to the presence of multiple different algal strains. Principal component analysis of the mycobiont data furthermore showed that gene expression differed significantly between lichen samples and mycobiont culture (Figure 4C).

We identified 1,749 differentially expressed mycobiont genes, of which 1,185 were upregulated in lichen thallus and 564 upregulated in mycobiont culture. Differentially-regulated genes (DEGs) were observed to lack functional annotation more frequently than across the entire transcriptome (Figure 4D). In total, 31% of the transcriptome failed to be assigned any function, as is typical for genome annotations of lichen fungi^10,22^, but among lichen-upregulated transcripts, this value reached 55%. Similarly, differentially expressed genes more often came from orthogroups identified as lichen-enriched (21% vs 11% in the whole transcriptome). We conclude that lichen-associated gene expression includes a large proportion of completely unknown gene functions.

**Figure 4.**
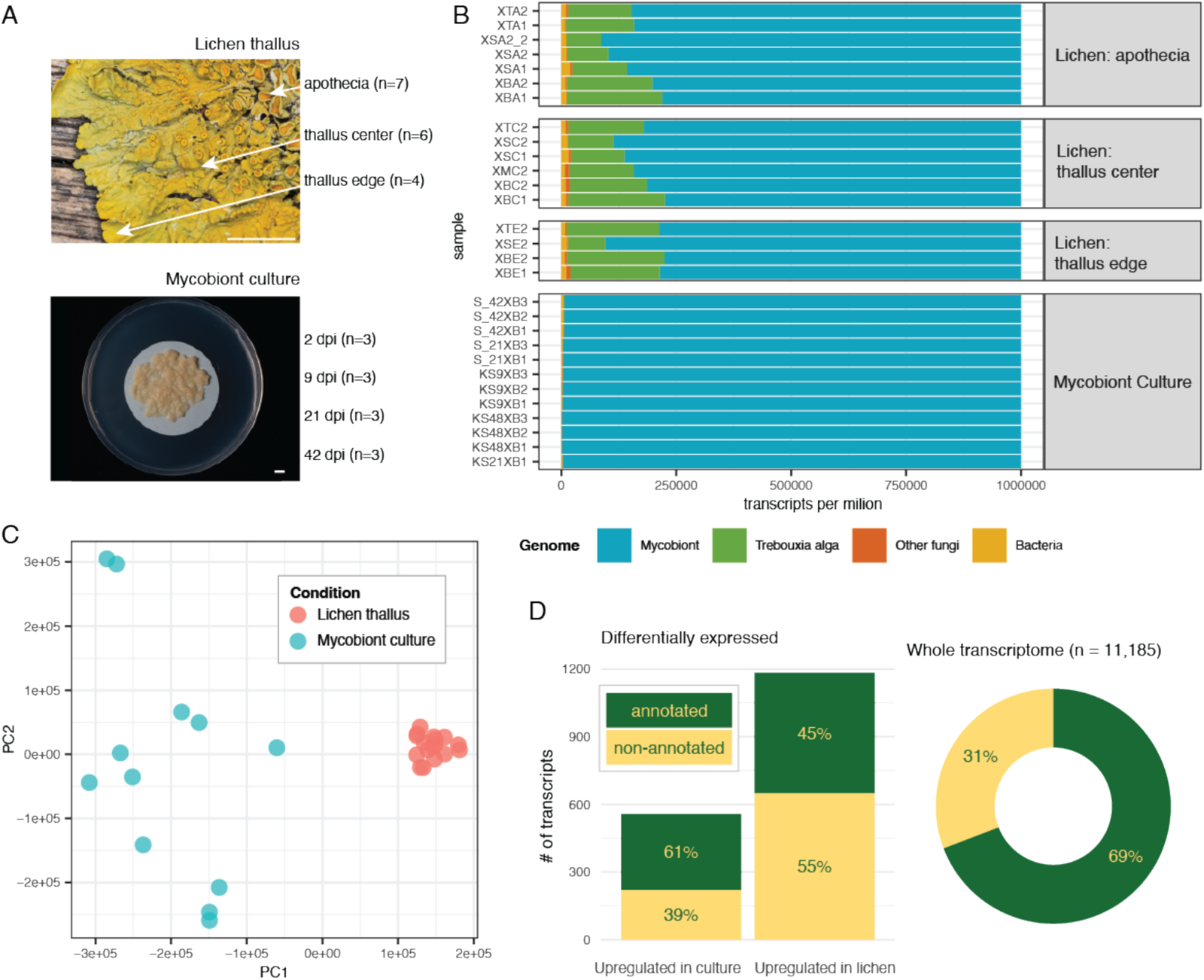
Transcriptomics of mycobiont culture and thalli of *X. parietina*. A. Samples used for RNA-seq: four timepoints of the fungus in culture (2 dpi, 9 dpi, 21 dpi, 42 dpi) and three developmental stages of lichen thallus (growing edge, center, and apothecia). Scale bars 5 mm. Apothecia are fruiting bodies formed by the fungus, but apothecia of *X. parietina* contain algal cells in the margin, and therefore the 7-22% share of algal transcripts is expected. B. Proportion of RNA-seq reads mapped to different categories of genomes. Transcript per million (TPM) values are summed across four groups: the mycobiont, *Trebouxia* algae, other fungi, and bacteria. C. Principal component analysis plot for RNA-seq samples colored by sample type. D. Proportion of differentially expressed transcripts with and without functional annotations (defined as any annotation with InterProScan or PFAM domains, or any assignment to UniProt, CAZy, MEROPS, Gene Ontology, and KEGG). Right panel shows percentage of transcripts with and without functional annotations across the entire transcriptome. See also Data S3.

### Genes related to cell division, cell wall biogenesis, secondary metabolism, and protein ubiquitination are upregulated in the lichen symbiosis

We next investigated the identity of gene functions differentially expressed in the lichen symbiosis compared to mycobiont culture. Transporters from the Major Facilitator Superfamily were overrepresented in both lichen thallus and mycobiont culture-upregulated genes (Figure 5A-B), with similar numbers of genes encoding transporters upregulated either in lichen thalli or mycobiont culture (Data S3; Figure 5C). However, genes encoding transporters believed to play an important role in the lichen symbiosis, such as putative polyol and ammonium transporters^1^, were lichen thallus-upregulated. Among nine genes highly similar to known polyol transporters, one (XANPAGTX0501_001653-T1) was upregulated in lichen thalli and also assigned to a lichen-enriched orthogroup. We also identified one lichen thallus-upregulated gene encoding an ammonium transporter (XANPAGTX0501_004972-T1). No genes encoding putative polyol and ammonium transporters were upregulated in mycobiont culture. Genes encoding proteins from other key functional groups implicated in fungal symbioses and/or fungal multicellularity^23^ such as transcription factors (TFs) and protein kinases were also differentially upregulated in either lichen thalli or mycobiont culture (Figure 5C). At the same time, specific groups of TFs showed patterns of differential expression. For example, homeobox domain TFs and Zinc finger C2H2-type TFs were upregulated only in lichen thalli. Similarly, three of four differentially expressed Zinc finger RING-type TFs were lichen thallus-upregulated. Representatives of these families of TFs have previously been linked to fruiting body development in mushroom-forming fungi^23^, consistent with a role in lichen tissue development. Conversely, the majority (six of eight) of differentially expressed Zn (II)2Cys6 zinc cluster TFs were upregulated in mycobiont culture.

In addition to 40 differentially expressed TFs, we identified other transcriptional regulators, upregulated in lichen thalli. Five genes encoding proteins with RNA-binding domains were upregulated in lichen thalli, for example, as well as one RNA-dependent RNA polymerase (XANPAGTX0501_002123-T1). More notably, a group of genes linked to protein ubiquitination were upregulated in lichen thalli (Figure 5C). These included eight genes encoding F-box proteins and four genes encoding BTB/POZ proteins (Data S3). Both these families are hypothesized to be involved in post-translational protein modification during formation of complex structures in mushroom development^23^, highlighting potential similarities in developmental biology of complex multicellular fungal structures.

Genes encoding proteins associated with cell division and growth– such as helicases, Rad21/Rec8-like proteins, and ribonucleases –were upregulated in lichen thalli (Figure 5C). Given the extremely slow growth of lichen fungi in culture^24^, this observation raises questions regarding whether the growth of mycobionts is conditional on the presence of other symbionts.

Genes associated with cell wall biosynthesis, cell wall remodeling, and cell wall proteins were often upregulated in lichen thalli (Figure 5C, Data S3). Genes assigned to Ricin B-like lectins, for instance, are overrepresented in lichen thallus-upregulated genes (Figure 5A). In addition, three genes with matches to Concanavalin A-like lectin/glucanase domains were lichen thallus-upregulated (Data S3). Four aspartic peptidases A1, involved in cell wall remodeling^23^, as well as seven carbohydrate-active enzymes (CAZymes) active on glucans and chitin were upregulated in lichen thalli.

The biosynthetic gene cluster (BGC) putatively responsible for biosynthesis of the anthraquinone parietin, the pigment of *Xanthoria*^25^ responsible for its yellow colour, was upregulated in lichen thalli (Figure 5D). Cluster Xp_GTX0501_17_Cluster_1 is a Type I polyketide BGC with similarity to the BGC of TAN-1612 (Data S1), a compound from *Aspergillus nidulans* structurally similar to anthraquinones^26,27^. Five more BGC were lichen thallus-upregulated, including Xp_GTX0501_4_Cluster_4, which is similar to a BGC linked to alkaloid peramine (Data S1). Such a BGC has been reported from the *X. parietina* mycobiont previously^27^, which reported a lack of A1 and R domains in the peramine synthase and deemed it non-functional. Based on our gene expression analysis, we can hypothesize that this BGC instead produces a different compound, that is induced during symbiosis. While only one BGC was upregulated in the mycobiont, some BGCs contained a mixture of thallus- and mycobiont-upregulated genes (Figure S3). Overall, somewhat contrary to expectation, genes related to secondary metabolism do not show a pattern of being lichen thallus-upregulated.

Differentially-expressed genes showed clear spatial clustering, suggesting that epigenetic regulation of gene expression might play a role in lichens (Data S3). Using a sliding window of 30 kbp, we therefore scanned the mycobiont genome to identify clusters of three or more jointly upregulated genes. We detected 92 lichen thallus-upregulated and 49 mycobiont culture-upregulated clusters. The majority (n=83 and 45 respectively) had fewer than 10 genes, however the largest cluster contained 19 lichen thallus-upregulated genes and was 87 kbp in length. Altogether, genes assigned to these clusters accounted for 47% of all differentially expressed genes. Interestingly, a similar pattern has been previously reported during fruiting body development in a non-lichenized ascomycete^28^.

**Figure 5.**
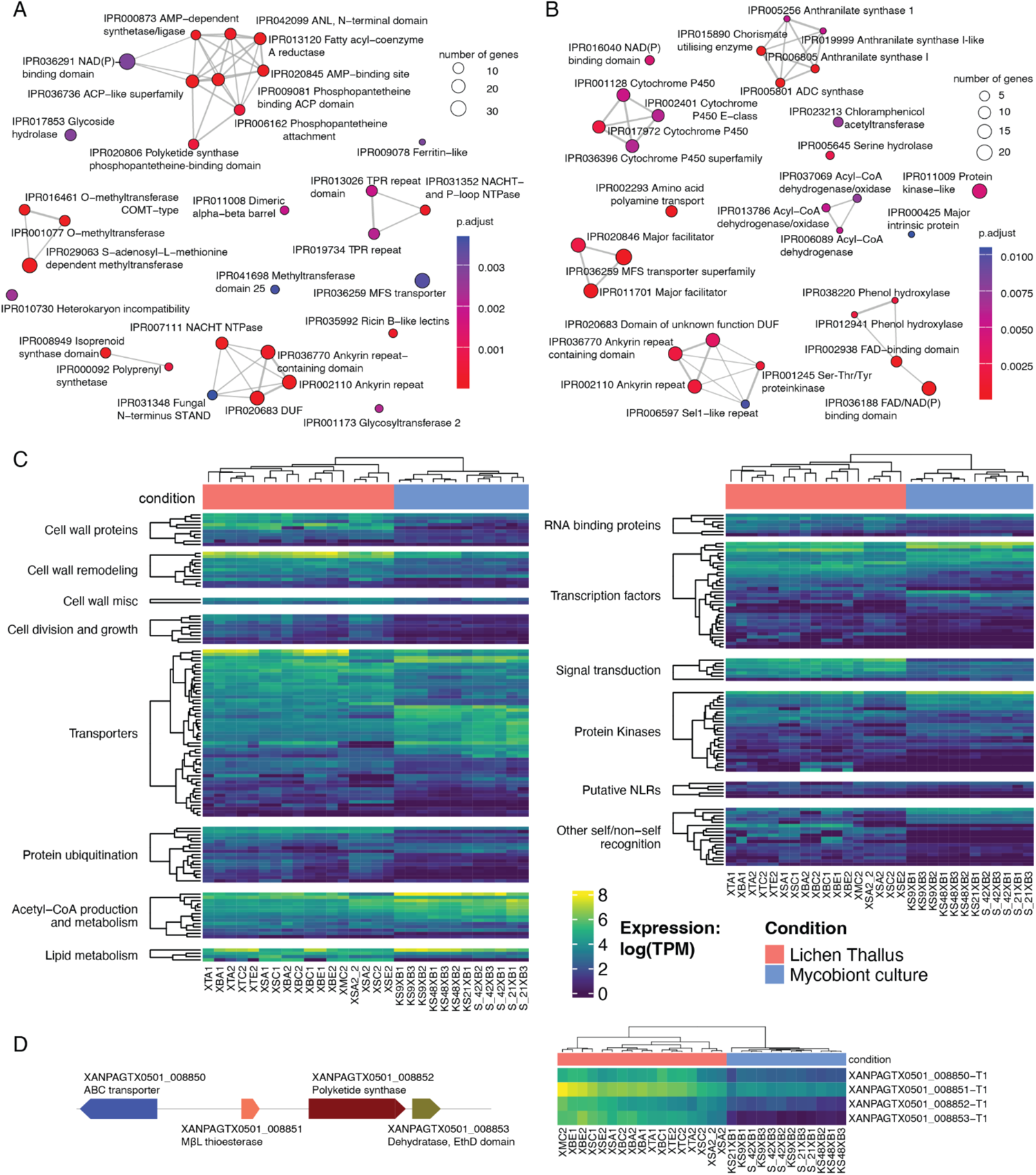
Differential Gene Expression of the mycobiont of X. parietina. A-B. Enrichment plots showing InterProScan domains enriched in genes upregulated in: A. Lichen thalli. B. Mycobiont culture. The size of the node represents the number of genes annotated with a given domain in the gene set. Two nodes are connected if domains are present together within at least one gene; the width of the edge corresponds to the number of such genes. C. Heatmap showing gene expression (as log(TPM), where TPM stands for transcripts per million). We only show differentially expressed genes assigned to one of the gene categories potentially involved in fungal multicellularity^23^ and symbiosis. Only categories with more than two genes are shown. D. Putative parietin biosynthetic gene cluster. The left track shows the structure of the cluster: ABC transporter, metallo-beta-lactamase-type thioesterase, polyketide synthase, and a dehydratase with an EthylD domain. The structure of Xp_GTX0501_17_Cluster_1 (Figure 5D) is identical to the anthraquinone BGCs identified in lichen genomes by Llewellyn et al.^26^. The heatmap shows expression levels for the four genes included in the cluster. See also Figure S3 and Figure S4.

### NLR-like genes are differentially expressed by *Xanthoria* mycobiont

We detected 23 genes in the mycobiont potentially encoding Nucleotide Oligomerization Domain (NOD)-like receptors, or NLRs – a group of proteins involved in self/non-self recognition and immunity in plants, animals, and fungi^29^ (Figure S4). These putative NLRs share structural features with known fungal NLRs^30^: a nucleotide-binding domain (either NB-ARC or NACHT), a repeat domain (either ankyrin, WD40, or tetratricopeptide repeats), and a variable effector domain. In half of putative NLRs, we identified effector domains, from three functional groups: enzymatic domains (alpha/beta hydrolase and nucleoside phosphorylase domains), cell death-inducing domains (HET and HeLo), and domains from InterProScan families IPR031359 and IPR031352 that lack a described function (Data S1). Unexpectedly, in the C-terminus of one putative NLR we found a papain-like protease domain that matched ubiquitin-specific proteases. The remaining 11 NLRs contained no conserved effector domain recognizable by InterProScan or PFAM– as is typical for fungal NLRs, whose effector domains are underrepresented in existing databases^30^.

Four putative NLRs are upregulated in lichen thalli, including one with a pore-forming HeLo domain (Figure 5C, Figure S4). Even though one NLR was identified as mycobiont culture-upregulated, its expression levels were low in only some of the lichen samples, while in others equivalent to expression in mycobiont culture samples. This inconsistency between lichen samples was more typical for genes potentially involved in self/non-self recognition (meaning NLRs and other genes with HET or HaLo domains), compared to other analyzed functional groups (Figure 5C). At the same time, of the 73 genes with HET or HaLo domains, 14 were consistently lichen thallus-upregulated. We conclude that a subset of NLRs may be associated with lichen development.

### Tissue-specific gene expression in lichen architectures

To investigate lichen morphogenesis, we carried out RNA-seq analysis of three distinct stages of lichen thallus development– the growing edge representing actively growing thallus, the center representing more mature thallus tissue, and apothecia which represent sites of sexual reproduction and ascospore formation (Figure 4A). We observed the largest number of up-regulated genes in apothecia (Figure 6A; Data S3). The highest number of DEGs was identified in a comparison of apothecia and growing thallus edge; while central thallus tissue was less differentiated from either tissue type. In the edge/center comparison, all but one center-upregulated gene was also among apothecia-upregulated genes (when we compared apothecia to the combined set of edge and center thallus tissue samples) (Figure S5). This pattern might be explained by apothecium primordia being present in the thallus center– in *X. parietina*, apothecia are clustered in the central part of the thallus^8^ –thereby affecting the expression profile making it more similar to that of the fruiting body, yet being too small to be detected and excluded during sample preparation. Our ability to detect tissue-specific patterns was also complicated by variation between lichen thalli, as expression profiles appeared to depend both on developmental stage and on each individual thallus preparation (Figure 6B). To identify DEGs between different developmental stages we therefore controlled for thallus identity, yet some tissue-specific genes may still have evaded detection.

Functional domains involved in protein ubiquitination and RNA interference were enriched among apothecium-upregulated genes (Figure 6C). The majority of lichen thallus-upregulated genes associated with these functions were also upregulated in apothecia compared to other developmental stages (Data S3, Figure S5). Most notably, XANPAGTX0501_008856-T1 encodes an F-box protein expressed in all lichen samples and none of the mycobiont culture samples. This gene had higher levels of expression in apothecia (b-value = 2.2 controlling for the thallus identity; Figure 6D). A similar pattern was observed in eight of 14 lichen thallus-upregulated genes encoding ubiquitination proteins, including four additional F-box proteins and all lichen thallus-upregulated genes associated with RNA interference (Figure 6D, Data S3). By contrast, the 27 lichen thallus-upregulated genes encoding transporters, and 23 lichen thallus-upregulated genes encoding transcriptional factors only included one and five apothecia-upregulated genes, respectively. Among the latter, we identified a homeobox protein gene (Figure S5), which are known to govern fruiting body formation in fungi^31^. Genes involved in karyogamy (*Kar5*) and conidiation (*Con6*) were also among genes upregulated in apothecia, as well as a gene encoding glycogen-debranching enzyme (Figure S5), consistent with gene expression profiles of mushroom development^23^. The gene model corresponding to the mating-type locus *MAT1-2-1* was also upregulated in apothecia, although the question remains whether *MAT* is actually functional since it contains a premature stop codon (Figure S6). While 71% (n=177 out of 250) of apothecia-upregulated genes were also upregulated in lichen thalli compared to mycobiont culture, 17 apothecia-upregulated genes were even more strongly expressed in culture. These genes included XANPAGTX0501_001643-T1, yet another apothecia-upregulated F-box gene. Similar to the lichen thallus / mycobiont culture comparison, the functions of most DEGs remain unknown. Of the 250 genes upregulated in apothecia compared to the other developmental stages, 171 (68%) have no functional annotations.

Contrary to our expectations, few genes were upregulated in the thallus edge compared to the center of the thallus. Since growth in *X. parietina* happens primarily at its narrow marginal rim, we expected numerous upregulated genes associated with active growth. However, no edge-upregulated gene was detected that could be linked to growth. One possible exception is XANPAGTX0501_009376-T1, which contains a ribosomal protein L10-like domain *(RPL10*; Figure 6E). While ribosomal proteins in general are associated with growth^23^, the profile of this gene does not match growth-associated genes discussed earlier. Instead of being lichen thallus-upregulated, it was more highly expressed in the mycobiont compared to any lichen sample. Alternatively, *RLP10* could be induced by stress, as is known for plant *RLP10*^32^, which is specifically expressed under UV light. Notably, another gene upregulated in the edge encodes the polyketide synthase linked to the biosynthesis of parietin – the key photoprotective pigment in *Xanthoria* (Figure 6E). Nearly half (n=6 out of 13) of edge-upregulated genes are predicted to encode secreted proteins, including a putative papain inhibitor (Figure 6E).

**Figure 6.**
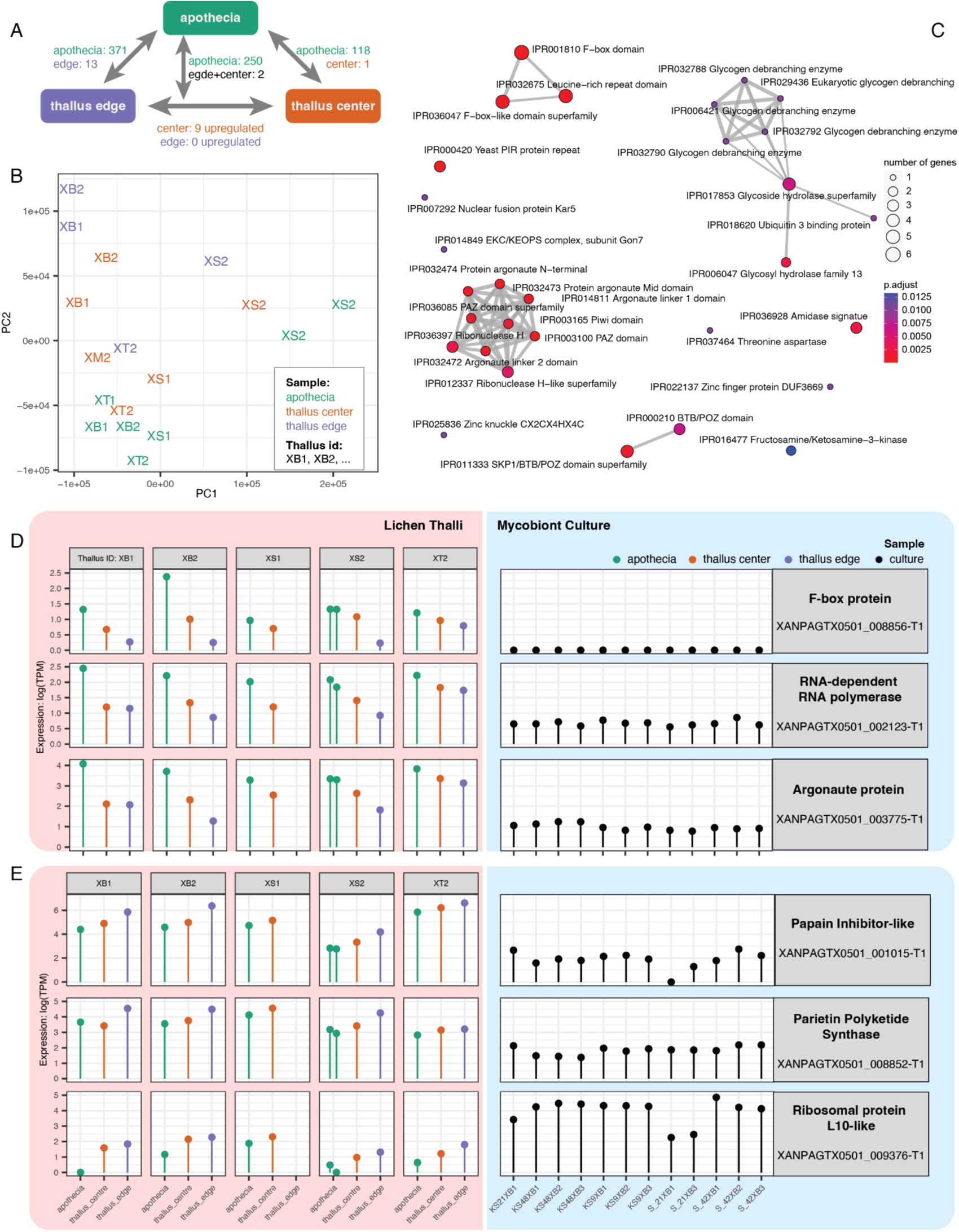
Differential Gene Expression of the mycobiont symbiont of *X. parietina* across three different developmental stages. A. Overview of differential gene expression between the stages. B. PCA plot for lichen-derived RNA-seq samples colored by the developmental stage. C. Enrichment plot showing InterProScan domains enriched in the apothecia-upregulated genes. D. Expression of three apothecia-upregulated genes involved in protein ubiquitination (F-box protein) and RNA interference (RNA-dependent RNA polymerase and Argonaute protein) across studied samples. The lichen samples are grouped based on the lichen thallus they derived from and colored based on developmental stage. The three shown genes were both upregulated in lichen samples compared to mycobiont culture, and in apothecia compared to other developmental stages. E. Expression of three thallus edge-upregulated genes. In addition to being upregulated in the thallus edge compared to apothecia, two of these genes are upregulated in lichen thalli compared to the mycobiont culture (putative polyketide synthase from the parietin gene cluster, and a putative papain inhibitor), and one is upregulated in the mycobiont culture (ribosomal protein L10-like). See also Figure S5 and Figure S6.

### The mycobiont secretome contains putative effector proteins

We next investigated whether mycobionts possess potential secreted effector proteins that potentially modulate cellular functions or impair immunity within symbiotic partners. Effectors are well known in pathogenic and mutualistic fungi ^33–36^. We identified 608 putative secreted proteins in the predicted proteome of *X. parietina* mycobiont, of which 154 were lichen thallus-upregulated and 40 mycobiont culture-upregulated (Figure S7A-B). As effectors are often sequence-unrelated, we carried out structural predictions using AlphaFold2 to identify structurally-related proteins within the predicted secretome. Structures with a quality score pTM≥0.5 (n=393) were used to construct a structural phylogenetic tree using FoldTree^37^. We divided the tree into 84 structural clusters, which together included 311 proteins (Figure 7A-B; Data S4); the remaining 82 proteins were considered singletons. In addition to structural analysis, we also screened the secretome using two effector-predicting tools: EffectorP and deepredeff (Figure S7C), although these provided inconsistent results.

The predicted secretome included proteins similar to known effectors. A large group of proteins (clusters 18-24a), for example, showed similarity to killer toxins Kp4 and a newly described effector from the plant pathogen *Zymoseptoria tritici* (Figure 7A,C). Collectively, these clusters accounted for 8% of the secretome and included 47 proteins, of which 18 were upregulated in lichen thalli compared to the mycobiont (Data S4). Protein XANPAGTX0501_009887-T1 was also lichen thallus-upregulated, and highly similar to Tsp1 (Figure 7D), an effector from *Trichoderma virens* that suppresses plant immunity by stimulating the salicylic acid pathway^38^. Other clusters of potential effectors include proteins with folds similar to known fungal effectors: CFEM proteins^39^, ribonucleases^40^, and NTF2/SnoaL proteins^41^ (Figure 7A). In the list of putative effectors, we also included thaumatin and gamma-crystallin-like proteins, as these families were identified as probable effectors^42,43^. Except for the ribonuclease cluster, these clusters had at least one lichen thallus-upregulated protein and none contained any mycobiont culture-upregulated proteins (Figure 7E). Similarly, of five proteins identified as Ricin B-like lectins, four were lichen thallus-upregulated, consistent with their proposed role in symbiont recognition^44^.

Secreted enzymes also account for over a third of the secretome (n=207) and are primarily represented by CAZymes and proteases. The most numerous enzyme cluster was formed by AA7 (Figure 7B) – oligosaccharide oxidase family expanded in lecanoromycete fungi^45^ and active on many substrates. Other major groups included GH16, a multifunctional family of glycoside hydrolases, and families active on beta-glucans (GH128, GH72, GH12), which might target the mycobiont’s own cell wall. Metallopeptidases M35 were also numerous, and curiously we identified several putative protease inhibitors (cluster 83), two of which were upregulated in lichen thalli. Unlike putative effectors, secreted enzymes were often upregulated in the mycobiont culture (n=19, out of 53 differentially expressed).

Combining sequence-based and structure-based annotation allowed us to assign putative functions to the majority of the secretome, although some assignments, especially based on the hits to the AlphaFold database, require significant further validation. However, the remaining 205 proteins contained no identified InterProScan or Pfam domain and yielded no significant match to a characterized protein when searched against structural databases (Data S4). These proteins might play a role in symbiosis, as the percentage of lichen thallus-upregulated proteins in the ‘novel’ set was even higher than in the secretome (33%, compared to 25% in the whole secretome and 11% across the whole transcriptome). The majority (n=165) of ‘novel’ proteins failed to produce structural models with quality scores above the set threshold (pTM≥0.5) and were consequently excluded from clustering. Others, however, were included and formed six clusters (cl04, cl14, cl15, cl48, cl52, cl82) composed entirely of proteins lacking annotation. Notably, cl04 consisted of eight proteins, two of which were differentially expressed and lichen thallus-upregulated (Figure 7A). Proteins from this cluster were classified as effectors by deepredeff, but not EffetorP. While all of them were assigned to lichen-enriched orthogroups, some of them showed similarity to uncharacterized proteins from various phytopathogenic and endophytic fungi, including *Alternaria alternata* and *Mollisia scopiformis* (Data S4), raising questions about their potential as novel lichen effectors.

**Figure 7.**
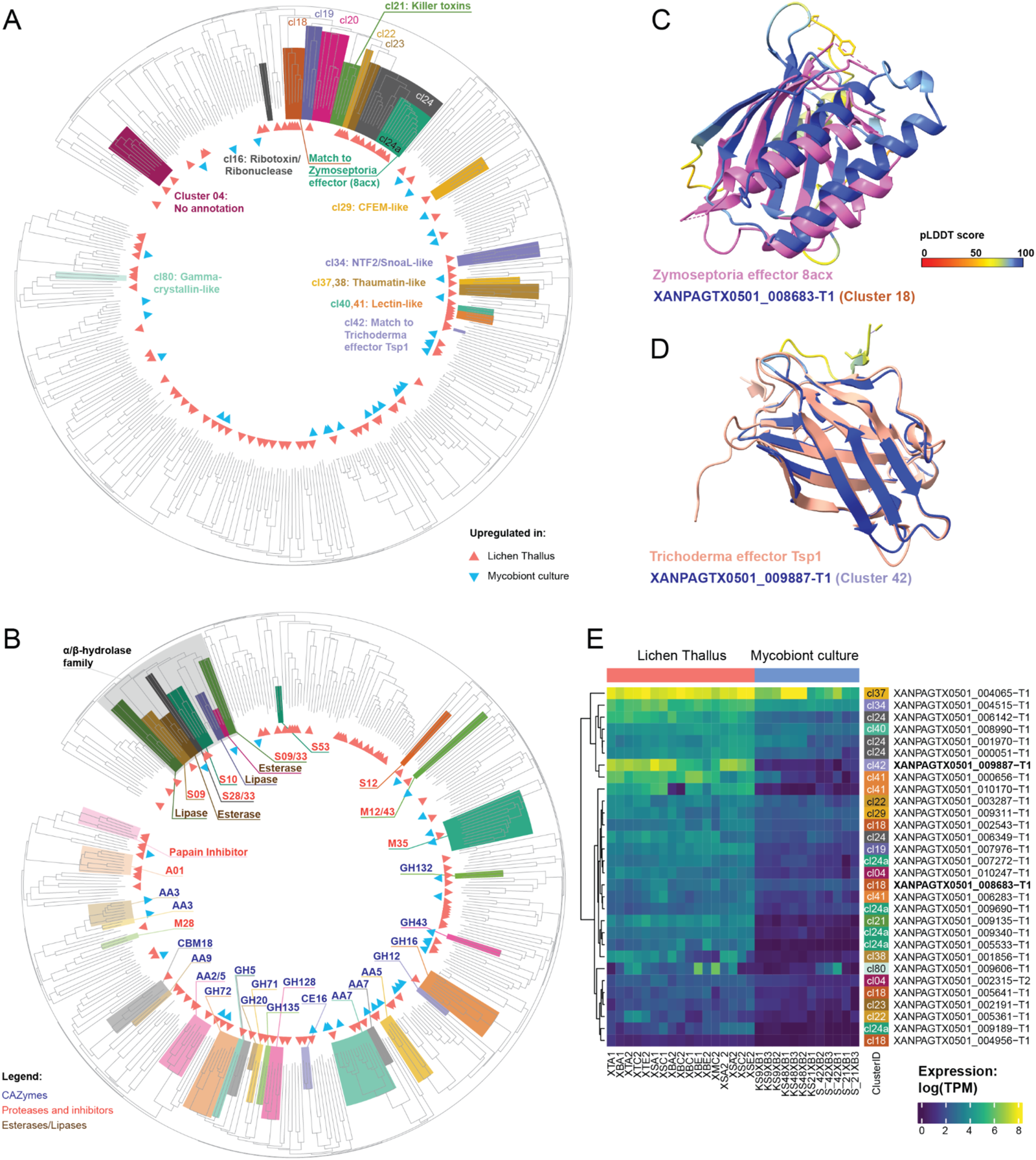
Structural clustering of the predicted secretome of the *X. parietina* mycobiont. A-B. The structural phylogenetic tree produced from structural models of predicted secreted proteins. Structural models with high confidence (pTM≥0.5) were analyzed using FoldTree, and the resulting tree split into 84 structural clusters (Data S4). The DEGs are indicated with triangles. A. Putative effector clusters and other clusters of interest are highlighted. B. Clusters formed by various hydrolases: Carbohydrate Active enZYmes (CAZymes), proteases, lipases, and esterases (as indicated by color). For the CAZymes and proteases, we give their family assignments. C. Predicted structure of XANPAGTX0501_008683-T1 (Cluster 18) superimposed onto a *Zymoseptoria* effector (PDB access number 8acx). D. Predicted structure of XANPAGTX0501_009887-T1 (Cluster 42) superimposed onto a *Trichoderma* effector Tsp1 (PDB access number 7cwj). E. Heatmap showing expression (as log(TPM), where TPM stands for transcripts per million) of DEGs corresponding to clusters of interest (shown in A). The right annotation track shows number of the cluster. The two proteins from C and D are highlighted in bold.

## Discussion

In this study we set out to explore how the intricate self-replicating architectures of lichens are formed from the symbiotic association of morphologically simple microorganisms. Our aim was to define the constituent microbiome of a single lichen species and to identify mechanisms that orchestrate lichen growth and development. To do this, we carried out a metagenomic analysis of a very common lichen *X. parietina*, investigated its development by transcriptional profiling, and used a combination of informatic and structural modelling to define potential determinants of lichen morphogenesis.

Our analysis of metagenomic data underlines the intrinsic complexity of lichen symbioses. While it has previously been assumed that one mycobiont individual corresponds to one thallus, our data suggest that multiple genotypes can be present within a single thallus. Non-standard MAF distributions in several of our samples (already reported from lichens before^46^) can potentially result from either the genotypes of male parents present in zygotes and ascospores within the fungal apothecia, or by several different mycobiont ‘individuals’ that live so closely together as to appear as one. Evidence for such ‘chimeric’ thalli have been reported previously^47,48^, and an experiment showed that a thallus fragment of *X. parietina* can fuse seamlessly back into its parent thallus^49^. Many of our samples also contained multiple lineages of the photobiont *Trebouxia*, as known for many lichens^50,51^, including *X. parietina*^52^. Multiple algal strains are hypothesized to offer benefit to the symbiosis by providing more plasticity, but may also reflect opportunistic acquisition of photobiont partners during lichen development.

Some lichen symbioses are now known to associate with certain bacteria and non-mycobiont fungi^3,4^. However, the organismal composition of *X. parietina* thalli was undescribed until now. Our metagenomic analysis reveals considerable diversity of microorganisms in addition to the mycobiont and the photobiont, including 157 distinct bacterial lineages. While basidiomycete yeasts, known from several lichen groups^3,53^, were not detected, ascomycete black yeasts were present occasionally. Of over a hundred bacterial species, the majority were also present sporadically or in samples collected from a specific substrate, while a smaller subset were present across all studied samples. A similar pattern was observed in the lung lichen *Lobaria*, where the bacterial microbiome can be split into a variable portion influenced by local environment, and a stable ‘core’ portion^54^. In *X. parietina*, the ‘core’ microbiota includes multiple lineages from the genus *Sphingomonas*, one of the bacteria most frequently detected in lichens^17^. Evidence from other lichen symbioses suggests that *Sphingomonas* is tightly associated with *Trebouxia* photobionts and can use polyols produced by *Trebouxia*^55,56^. Whether *Sphingomonas* and other bacteria are commensals or mutualistic symbionts in *X. parietina* remains to be tested, however their presence needs to be taken into account both in terms of our understanding of lichen biology, and also during the experimental design and analysis of lichen-derived ‘omics’ data.

Using transcriptional profiling, we next identified biological processes that differ between the mycobiont in its natural state– as a member of the complex lichen symbiosis– and in aposymbiotic pure culture. As in previous reports^57–59^, polyol and ammonium transporters were upregulated in lichen thalli, consistent with the hypothesis that mycobionts use alga-produced polyols in exchange for ammonium. Cell wall proteins, including lectins, were also upregulated in symbiosis, matching a predicted role in hyphal adhesion and symbiont recognition^44^. Another class of lichen thallus-upregulated cell wall protein, fungal hydrophobins, have also been shown to be differentially expressed in *X. parietina* and hypothesized to play a role in creating lichen architectures^60^. Secondary metabolism, cell wall synthesis and remodeling, and cell division functions are also associated with lichen thalli.

We were particularly interested in defining how lichen gene expression is regulated during development. It appears that changes of gene expression during lichenisation may be partially driven by epigenetic regulation, because differentially expressed genes tend to form clusters instead of being randomly dispersed across the fungal genome. Among identified differentially expressed transcription factors (TFs), we identified different classes of TFs being upregulated in different states, consistent with orchestration of a specific developmental programme, while post-transcriptional and post-translational regulation also probably play an important role, as we identified lichen thallus-upregulated genes potentially involved in RNA-interference and targeted protein ubiquitination. Among the latter, F-box and BTB/POZ proteins, which have been linked to complex multicellularity in mushroom-forming fungi and non-lichenized ascomycetes^23,61^, were especially prevalent. Surprisingly, however, we did not identify many differentially-expressed genes among key signaling pathways such as MAPK-dependent signaling and TOR signaling.

Studies of complex multicellularity in fungi primarily focus on fruiting bodies^23^, which have anatomy similar to lichens (as both are essentially formed by tightly packed hyphae), but which also differ from lichens in two important ways. First, lichen thalli are formed through concerted growth of multiple symbionts. Second, in mushroom-forming fungi, complex architectures emerge only to serve a specific function– usually sexual reproduction although not exclusively as in for instance sclerotia –and the fungus exists primarily as simple mycelium. For mycobionts, however, complex lichen thallus architectures represent the *only* known mode of existence, and lichen thalli are not primarily linked to reproduction. In many lichens, for example, none of the symbionts undergo sexual reproduction. Thalli of *X. parietina* contain apothecia of its mycobiont, yet most of the body is formed by vegetative hyphae and photobiont cells. This prompted us to try to separate the mechanisms behind complex structures from those specific to fruiting body development and sexual reproduction. We identified genes upregulated in apothecia compared to vegetative parts of the thallus. In addition to expected genes involved in sexual reproduction, we identified numerous genes linked to RNA-interference and targeted protein ubiquitination, consistent with a role in fruiting body development. The majority of apothecium-upregulated genes were also upregulated in lichen thalli compared to cultures (as expected, given that mycobionts never reproduce sexually in culture^2^). Several exceptions– genes upregulated in fruiting bodies *and* the mycobiont culture –are of special interest with regard to a role in fungal development. Also of interest are genes upregulated in lichen thalli compared to culture but *not* in apothecia, as these might represent core machinery required for forming complex lichen structures. These include the majority of lichen thallus-upregulated cell wall proteins and TFs, as well as genes linked to self/non-self recognition (discussed below). When considered together, our study provides evidence that a similar toolbox is used for complex multicellularity by lichen mycobionts and non-lichenized ascomycete and basidiomycete fungi, consistent with a recent hypothesis^62^, and suggesting conservation in higher order fungal developmental biology.

The longevity of lichen thalli is another feature separating them from complex fungal fruiting bodies. *X. parietina* and similar lichens grow primarily in the thin outer rim of the thallus, meaning that lichen tissue is younger at the margins of a thallus. We therefore used different parts of thalli as a proxy for lichen developmental stages. We aimed to identify genes involved in active growth of a lichen thallus and tissue differentiation. However, our analysis yielded only a few thallus edge-upregulated genes. The lack of growth-associated gene expression in the edge might be explained by both biological and technical reasons. While growth in *X. parietina* happens primarily at its narrow marginal rim, central parts of thalli are also capable of regenerative growth^8^. In addition, lichen growth does not occur continuously and instead switches on and off depending on microclimate^63^. Both factors might therefore complicate detection of growth-related genes in a transcriptomic study. Genes upregulated in the thallus edge include several secreted proteins and a gene cluster linked to pigment biosynthesis. This can be seen as evidence that lichen tissue in this developmental stage experiences stress and secretes proteins in order to affect other microorganisms or modify its substrate. Our study is the first to compare gene expression profiles of different parts of a lichen architecture and has allowed broad classification of lichen-associated gene functions by developmental stage.

Our final aim was to investigate factors that mediate the interaction between a mycobiont and other symbionts. The lichen symbiosis likely involves bidirectional recognition between symbionts and potentially the recruitment of appropriate strains. While much is known about fungal self/non-self recognition systems, research mostly focuses on the mechanisms for different strains within one species to recognize each other, and our understanding of how fungi recognize other organisms remains poor^64^. NLRs and HET domain proteins are hypothesized to play a role in fungal immunity and fungal symbioses, but experimental validation is still lacking^64^. We identified putative NLR-encoding genes in the *X. parietina* mycobiont, several of which were upregulated in lichen thalli, suggesting a role in lichen development or maintenance. We should note, however, that our data derived from established lichen thalli, and therefore we were unable to capture gene expression changes during initial establishment of the symbiosis. Future research will need to test the role of NLRs in recognition between lichen symbionts.

In addition to recognizing each other, symbionts may possess machinery to influence one another during lichen formation. Secreted effector proteins are often used by fungi, both pathogenic and mutualistic, to establish contact and suppress the immune response of their plant host^33–36^. Since green algae share some features of the plant immune system^65^, we recently hypothesized^66^ that mycobiont effectors might play a role in lichen symbioses. By predicting and analyzing the secretome of the mycobiont, we identified putative effectors, many of which were upregulated in lichen thalli. Effectors often evolve so rapidly that they lose sequence similarity^67^, so we predicted protein structures for all 608 secreted proteins and used structures to group similar secreted proteins together. In this way, we were able to identify a large group of proteins with similarity to killer toxin Kp4 and several fungal effectors^68,69^. We conclude that effector-like proteins are encoded by the mycobiont, consistent with manipulation of other symbionts during lichen development. In addition, our analysis revealed completely novel proteins that show no similarity to any characterized protein that might represent novel lichen-specific families of effectors.

## Conclusion

In summary, metagenomic and metatranscriptomic analysis of *X. parietina* has identified biological processes involved in lichen development. Our results show that lichen morphogenesis shares features with development of multicellular structures by non-lichenized fungi, such as sclerotia and mushrooms, but it is also clear that lichen formation involves a large amount of unknown biology. The majority of genes upregulated in the symbiotic state cannot, for example, assigned any function based on similarity to databases. We attempted to push this boundary by making structural predictions for proteins secreted by the mycobiont, which allowed identification of structurally-related putative effectors, but also highlighted the large number of completely novel proteins present in lichens. Our study will therefore provide a resource for future research on developmental biology of this elusive group of organisms.

## Acknowledgments

This work was supported by grants from the Leverhulme Trust RPG-2018-139, The Gatsby Charitable Foundation, the Halpin Family and the Biotechnology and Biological Sciences Research Council BBS/E/J/000PR9798 to NJT. We thank Jesper Svedberg and Markus Hiltunen Thorén for suggestions on data analysis, Paul Dyer for providing the mycobiont culture, Phil Robinson for providing photographs, and Alison MacFadyen for help with depositing data.

## Author contributions

NJT, KS, and GT conceived the project. KS, XY, and GT performed lab work. GT, NS, CJ, DM, AB, ARB, SLAV, IOB, HJ designed and performed bioinformatic analysis. GT prepared figures and tables. GT and NJT drafted the manuscript, and all authors contributed to editing.

## Star Methods

### RESOURCE AVAILABILITY

#### Lead Contact

Nicholas J. Talbot is the lead contact for this study.

#### Materials Availability

*X. parietina* lichen samples generated in this study will be made available upon request from the lead contact.

#### Data and Code Availability

● Raw metagenomic and metatranscriptomic sequencing data, as well as assembled and annotated genomes have been deposited at ENA (PRJEB78723 and PRJEB38537; data release pending).
● Code generated for data analysis is available in GitHub (https://github.com/metalichen/2024-Multipartite-complexity-omics-Xanthoria).

### EXPERIMENTAL MODEL AND SUBJECT DETAILS

#### Lichen thalli

Lichen thalli were collected in Norwich Research Park (Norwich, UK; 52.623133°N, 1.221621°E). For the metagenomes, eight thalli of *X. parietina* were collected from tree bark and concrete (Data S5). Three were incubated in a growth chamber for 12 months under a 12-h night/day light cycle. These thalli were sprayed weekly alternating between deionized water and liquid Bold’s Mineral Medium (BMM). The rest of the thalli were sourced from the field and immediately used for DNA extraction following air drying. One additional thallus of *X. calcicola* was collected from concrete substrate. For the metatranscriptomes, seven thalli were collected from tree bark, tree twigs, concrete, and metal substrates (Data S5).

#### Xanthoria parietina mycobiont culture

A pure culture of *X. parietina* mycobiont was kindly provided by Prof. Paul Dyer, University of Nottingham, UK. The culture was obtained from a thallus collected in the Peak District, UK. The culture was maintained in liquid Malt Extract Yeast Extract (MEYE) medium.

### METHOD DETAILS

#### Mycobiont genome sequencing and assembly

A fragment of a *X. parietina* thallus was cleared from all visible contaminants and all apothecia removed with a razor blade. Lichen material was homogenized with a Geno/Grinder homogenizer (SPEX SamplePrep, Metuchen NJ, USA) at 1300 rpm for 1 min. DNA was extracted with the NucleoBond High Molecular Weight DNA Kit (Macherey–Nagel, Düren, Germany). Short fragments were removed with Circulomics Short Read Eliminator Kit (Pacific Biosciences, Menlo Park CA, USA) with 25 kbp cut-off. The resulting 0.6 μg of high-molecular weight DNA were used for long-read sequencing. The library was prepared using a DNA ligation V14 kit (Oxford Nanopore Technologies, Oxford, UK) and sequenced using a PromethION Flow Cell FLO-PRO114M (Oxford Nanopore Technologies, Oxford, UK). We used Dorado v0.2.4 (Oxford Nanopore Technologies, Oxford, UK) for base-calling. Contigs were assembled with Flye v2.9-b1780 with ‘overlap 10 K, error rate 0.01, no-alt-contigs, meta’ flags. Long-read sequencing and assembly were performed by Future Genomics (Leiden, Netherlands).

#### RNA extraction and sequencing

For transcriptomes of the mycobiont culture, the culture was plated on nitrocellulose filters incubated on 2% agar plates with BMM:MEYE 99:1 medium (following Joneson et al.^71^). The cultures were harvested at 2, 9, 21, and 42 days post inoculation; each time point had three replicates. The cultures were snap-frozen in liquid nitrogen and RNA was extracted using an RNeasy Plant Mini Kit (QIAGENE, Hilden, Germany). Total RNA was sent to Novogene UK (Cambridge, UK) and sequenced on an Illumina HiSeq2500 platform to PE150 data.

For lichen metatranscriptomes, we air dried collected samples and separated each into three developmental stages (central part, thallus edge, and apothecia) manually with a razor blade. RNA was extracted and sequenced as described above. From seven thalli, we produced 17 metatranscriptomes, of which six were derived from the central part, four for the thallus edge, and seven from apothecia (Data S5).

#### Mycobiont genome annotation

First, we removed non-mycobiont sequences from the assembly, by using a metagenomic binning approach. We used short-read data produced from the same lichen sample as the long-read assembly (Data S5), and this was aligned to the long-read assembly with bowtie2^72^. Using the resulting alignment, we binned the assembly with MetaBAT2^73^. To identify the bins corresponding to the mycobiont genome, we used EukCC v2.1.2^74^. We also ran a BLASTx search against the NCBI-nr database using each contig as a query. The final MAG contained 58 contigs and was created by combining two bins and three unbinned contigs with hits to Lecanoromycete fungi. The quality of the MAG was assessed with EukCC and BUSCO5^75^ using the ascomycota_odb10 database. The mitochondrial genome was detected using the same BLASTx search. To identify telomeric repeats, we used a script from Hiltunen et al.^76^ with TAA[C]+ as a query.

Prior to gene annotation, we masked repeat elements in the genome. We created a custom repeat library using RepeatModeler v2.0.3^77^ with the -LTRStruct flag. Using the repeat library, we masked repeats in the genome using stand-alone RepeatMasker v4.1.2 (https://www.repeatmasker.org/). To annotate repeat-induced point mutations we used RIPper^78^. Gene prediction and functional annotation was done with the Funannotate pipeline v1.8.15^79^.

Gene prediction parameters were generated using the ‘funannotate train’ module with the transcriptomic data from the mycobiont culture as an input. For gene prediction, we used the ‘funannotate predict’ module, which performed *ab initio* prediction with Genemark-ES v4.62^80^, Augustus v3.3.2^81^, CodingQuarry v2.0^82^, GlimmerHMM v3.0.4^83^, and SNAP 2006-07-28^84^. We created consensus models with EVidence Modeler v1.1.1^85^ and annotated tRNA with tRNAscan-SE v2.0.9^86^. To create functional annotations, we used the ‘funannotate annotate’ module, which runs HMMER v3.3.2 and diamond v2.1.6^87^ searches against the following databases: PFAM v35.0^88^, UniProtDB v2023_01^89^, MEROPS v12.0^90^, dbCAN v11.0^91^, and BUSCO ascomycota_odb10^75^. In addition, we annotated the predicted proteins using Emapper v2.1.12^92^ and Eggnog v5.0 database, InterProScan v5.42-78.0^93^, antiSMASH v7.0 web server^94^, and KAAS web server^95^. To further improve the gene annotation prediction, we employed a sequence homology-based approach. We used the orthogroup clustering method (see Identify lichen-enriched orthogroups below), focusing only on *X. parietina* genes. From these, we leveraged the previously identified functional annotations from Funannotate and assigned gene functions to orthogroups. If at least 40% of the genes within an orthogroup were annotated in *X. parietina*, we assigned the remaining *Xanthoria* genes the same functional annotation.

To predict the secretome, we used three tools: WolfPSORT^96^, deepTMHMM^97^, and SignalP v5^98^. We defined a protein as being putatively secreted using three criteria: signal peptide identified by SignalP, no transmembrane domains identified by deepTMHMM, and the probability of being secreted of ≥0.6 according to WolfPSORT. All secreted proteins were analyzed with two effector-predicting tools: EffectorP v3.0 ^99^and deepredeff v.01.1^100^. To identify NLR-like proteins, we used a custom script filtering proteins based on their InterProScan domains (Data S1); the list of domains typical for fungal NLRs we took from Uehling et al^29^; the visualization script was partially based on RefPlantNLR^101^. To annotate the *MAT* locus, we ran a BLASTp search against the predicted proteome. As a query, we used *MAT* genes from *X. polycarpa* (GenBank IDs: CAI59767.1, CAI59768.1, CAI59769.1, CAI59770.1, CAI59771.1, CAI59772.1). We used the same queries to screen metagenomic assemblies and raw reads (see below). To identify putative polyol transporters, we used a BLASTp search with four known transporters (GenBank IDs: AAX98668.1, CAR65543.1, CAG86001.1, NP_010036.1) as queries; hits with the e-value<1e-100 were considered.

We annotated the mitochondrial genome using MitoFinder v1.4.1^102^. As a reference, we used the mitochondrial genome of *Peltigera malacea* from Xavier et al^103^. We added the missing *rrnL* annotation manually based on the BLASTn search results.

#### Identification of lichen-enriched orthogroups

To identify orthogroups enriched in lichen-forming fungi, we analyzed a dataset of 44 fungal species, including 18 lichen-forming and 26 non-lichen-forming fungi (Data S2). We employed Orthofinder v2.5.4^104^ to classify proteins from these species into orthogroups. The copy number matrix from these orthogroups, was then subjected to the fisher.test function in R to identify orthogroups that have an overrepresentation of genes present predominantly in lichen-forming fungi when compared to fungi that do not form lichens. This function uses an ABCD matrix to calculate the enrichment, where A represents the total number of genes in a specific orthogroup in lichen-forming fungi, B represents the total number of genes in the same orthogroup among non-lichen forming fungi, C represents the total number of genes in remaining orthogroups in lichen-forming fungi and D represents the total number of genes in remaining orthogroups among non-lichen-forming fungi. The orthogroups significantly enriched with lichen genes were ones with a Benjamin-Hochberg corrected p-value ≤0.05.

#### Metagenome sequencing and analysis

Eight samples of *X. parietina* and one of *X. calcicola* were collected and air dried (Data S5). The samples were homogenized as described above and DNA was extracted with a DNeasy Plant Mini Kit (QIAGENE, Hilden, Germany). DNA was sequenced on an Illumina NovaSeq 6000 platform by Novogene UK (Cambridge, UK).

Metagenomic data from *X. parietina* samples were cleared from human contamination by aligning to the reference human genome with bowtie2^72^. We removed adapters using cutadapt v1.17^105^. The filtered data were assembled using MEGAHIT v1.2.6^106^. We ran both individual assemblies for each sample, and co-assembly of all *X. parietina* samples. Next, all assemblies were binned with MetaBAT v2.15^73^. To identify eukaryotic MAGs and assign them preliminary taxonomic assignments, we screened all bins with EukCC2^74^. For prokaryotic MAGs, we used CheckM v1.2.0^107^. Next, we selected all high and medium quality MAGs (completeness ≥50%, contamination <10%)^108^ and dereplicated them using dRep v2.5.0^109^ at 95% ANI (average nucleotide identity) and 40% AF (alignment fraction) thresholds in order to obtain species-level representatives.

To produce taxonomic assignments for the eukaryotic MAGs, we combined them with reference genomes (Data S2) and built phylogenomic trees. The MAGs were split into two groups – fungal and algal – based on the annotations from EukCC. To the fungal tree we also added the long-read assembly of the mycobiont. For the two reference algal genomes that lacked annotations (Data S2), we ran BUSCO5 with the chlrophyta_odb10 database and used the predicted proteins for the analysis. The species tree was generated using OrthoFinder v2.5.4. The MAG of the mycobiont was identified based on its position in the phylogenomic tree. To confirm this, we aligned it against the long-read genome assembly of *X. parietina* mycobiont using Minimap2 v2.24-41122^110^.

To assign taxonomy to the bacterial MAGs, we used GTDB-Tk v1.7.0^111^ with the GTDB database v202^112^. From the alignment of 120 marker genes produced by GTDB-Tk, we generated a maximum-likelihood phylogeny using IQ-TREE v2.2.2.2^113^.

To map the presence/absences of species-level lineages across the metagenomic samples, we used the metaMap pipeline (https://github.com/alexmsalmeida/metamap). Reads from all metagenomes, including the additional *X. calcicola* sample, were aligned against the entire MAG catalog with BWA v0.7.17-r1188^114^. Secondary alignments were removed using Samtools v1.10^115^. All MAGs covered ≥50% in a given metagenome were counted as present. To calculate the depth of coverage, we multiplied the number of reads aligned to the MAG by the read length and divided by the total length of the MAG.

For metatranscriptomic analysis, all MAGs except for the mycobiont MAG, were annotated. First we filtered each MAG using Funannotate modules clean and sort to remove contigs shorter than 500 bp and showing >95% overlap with other contigs. We masked repeats in the eukaryotic MAGs using RepeatMasker and the RepBase database v18.08^116^. For fungal MAGs we used fngrep.ref, which contains repeats from across fungi; for algal MAGs we used chlrep.ref, which contains annotated repeats from *Chlamydomonas*. Next, we ran the ‘funannotate predict’ module as described above. For training Augustus, we used the BUSCO dikarya_odb9 database for fungi and chlorophyta_odb10 for algae. Bacterial genomes were annotated using Prokka v1.14.6^117^.

#### Confirming sample identities

To confirm the identity of mycobionts from metagenomic and genomic samples, we ran a BLASTn search to extract the ITS region (ITS1, 5.8S ribosomal RNA gene, ITS2) using JF831902.1 *X. parietina* as the query. We combined the extracted ITS sequences with 338 reference sequences from various Teloschistaceae (Data S5) and aligned them using MAFFT v7.271^118^ with the –maxiterate 1000 flag. The alignment was clipped using trimAL v1.2^119^ to remove positions present in <70% of sequences. The phylogeny was calculated using IQ-TREE^113^.

#### Ploidy analysis

To calculate the minor allele frequency distributions of the mycobiont genome, we adapted the pipeline from Ament-Velásquez et al^120^. We aligned the metagenomic short-read data to the long-read mycobiont genome assembly using BWA v0.7.17-r1188^114^ with PCR duplicated marked with Picard v2.21.2 (https://broadinstitute.github.io/picard/). We called variants using Varscan v2.3.9^121^ with the flags --p-value 0.1 --min-var-freq 0.005. We removed contigs shorter than 100 kpb and filtered out variants overlapping with repeat elements. The resulting vcf file was processed using the vcfR library v1.15.0^122^.

#### Transcriptomic analysis

We trimmed the data to remove adaptors and poly-A tails with cutadapt^105^. To remove rRNA contamination, we used SortMeRNA v3.0.3^123^ using the Silva database v132^124^. Next, we created a reference index by combining predicted coding sequences from the annotated MAGs and the long-read mycobiont genome. We pseudoaligned the transcriptomic data to the index using kallisto v0.46.2^125^. For differential gene expression analysis of the mycobiont, we used sleuth v0.30.1^126^. Genes were identified as differentially expressed if they had |b-value| >1 and P-adjust < 0.05. To compare samples from different developmental stages, we controlled for the thallus identity following https://pachterlab.github.io/sleuth_walkthroughs/pval_agg/analysis.html. For enrichment analysis, we used ClusterProfiler v4.2.2^127^. To identify clusters of differentially-expressed genes, we used CROC^128^

#### Protein structure prediction and analysis

We predicted structures of the proteins from the predicted secretome using ColabFold v1.5.0^129^. We used FoldSeek v8.ef4e960^130^ to search the structures against two databases: PDB^131^ (downloaded on 2023.12.11) and AlphaFold^132^ (downloaded on 2024.04.18). We only retained the hits with e-value <0.001. All protein structures with pTM (template modeling score) ≥0.5 were subjected to structural clustering. We used the 0.5 threshold following Seong and Krasileva^67^. For cultering, we first removed the signal peptide (as identified by SignalP, see above) and disordered regions, defined as residues with pLDDT (predicted local-distance difference score) ≤0.55. Next, we constructed a structural phylogenetic tree using FoldTree^37^. Based on the LDDT tree produced by FoldTree, we manually curated a set of clusters with similar protein structures. One cluster (cl42) included only one protein. It was designated cluster status due to its similarity to a known effector. To visualize the structural tree, we used iTOL v6^133^. The protein models were visualized using ChimeraX v1.6.1^134^.

