## Supplementary material for "Multipartite complexity of the lichen symbiosis revealed by metagenome and transcriptome analysis of *Xanthoria parietina*": Figures S1-S7

**Supplementary Figures**

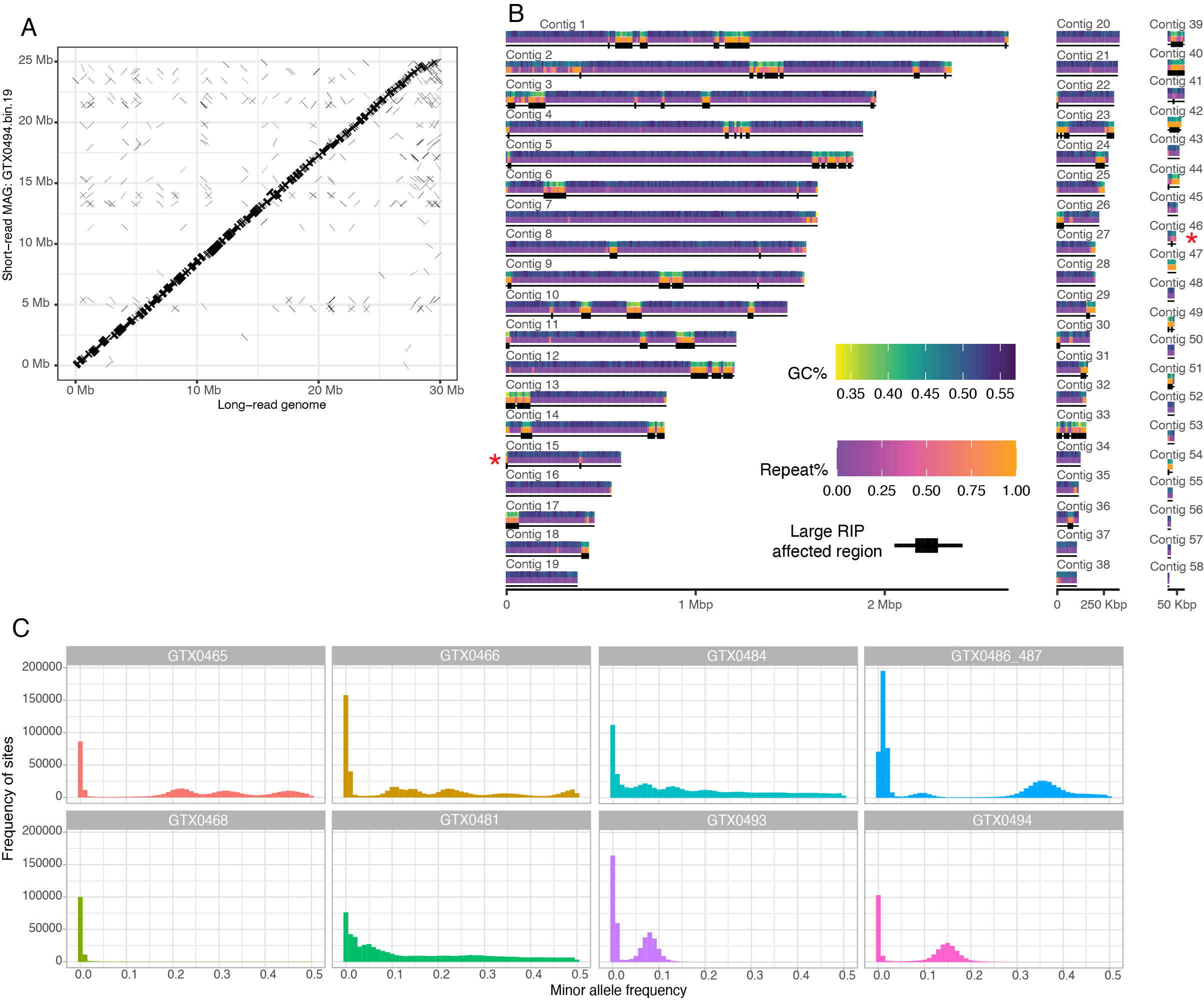

**Figure S1. Genome of X. parietina mycobiont in metagenomic data.** A. Whole genome alignment of the long-read genome assembly of *X. parietina* mycobiont and the metagenome-assembled genome of the same fungus extracted from the metagenomic data. The alignment was generated using Minimap2 v2.24-41122. B. *X. parietina* mycobiont nuclear genome. Each contig is represented by three annotation tracks: GC content, repeat content, and presence of Large RIP Affected Regions (LRARs). The x-axis corresponds to contig length. Red asterixis show telomeric repeats. Contig labels are shown next to them. C. Minor allele frequency plot for the *X. parietina* mycobiont. We aligned metagenomic reads against the newly produced genome of the *X. parietina* mycobiont.

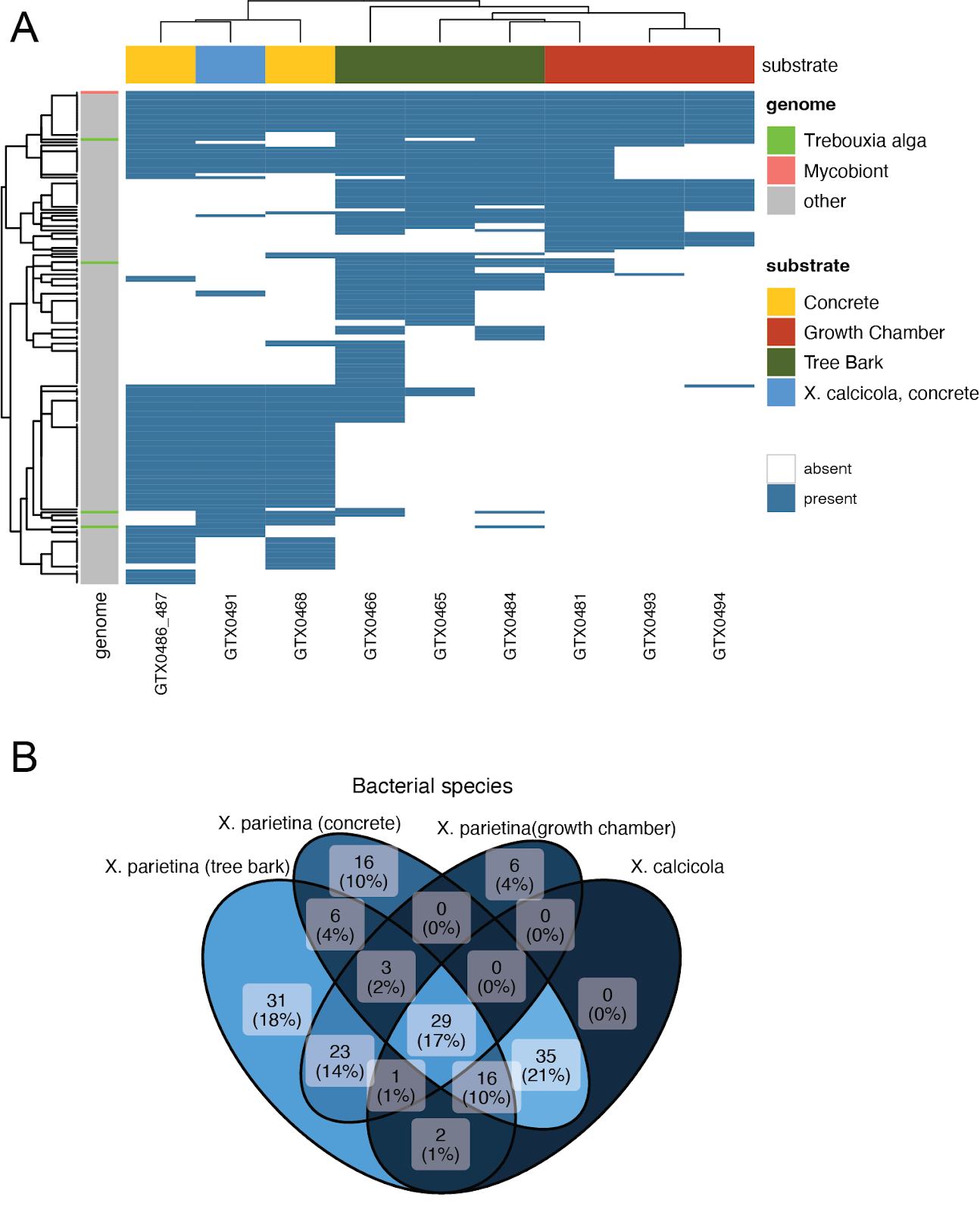

**Figure S2. Comparing species composition of *X. parietina* to a *X. calcicola* sample.**  A. Presence/absence map of 168 *X. parietina*-derived MAGs in nine metagenomes: eight from *X. parietina* and one sample of *X. calcicola*. B. Venn diagram comparing the lists of bacterial species-level lineages in *X. calcicola* and *X. parietina* (the latter split by substrate).

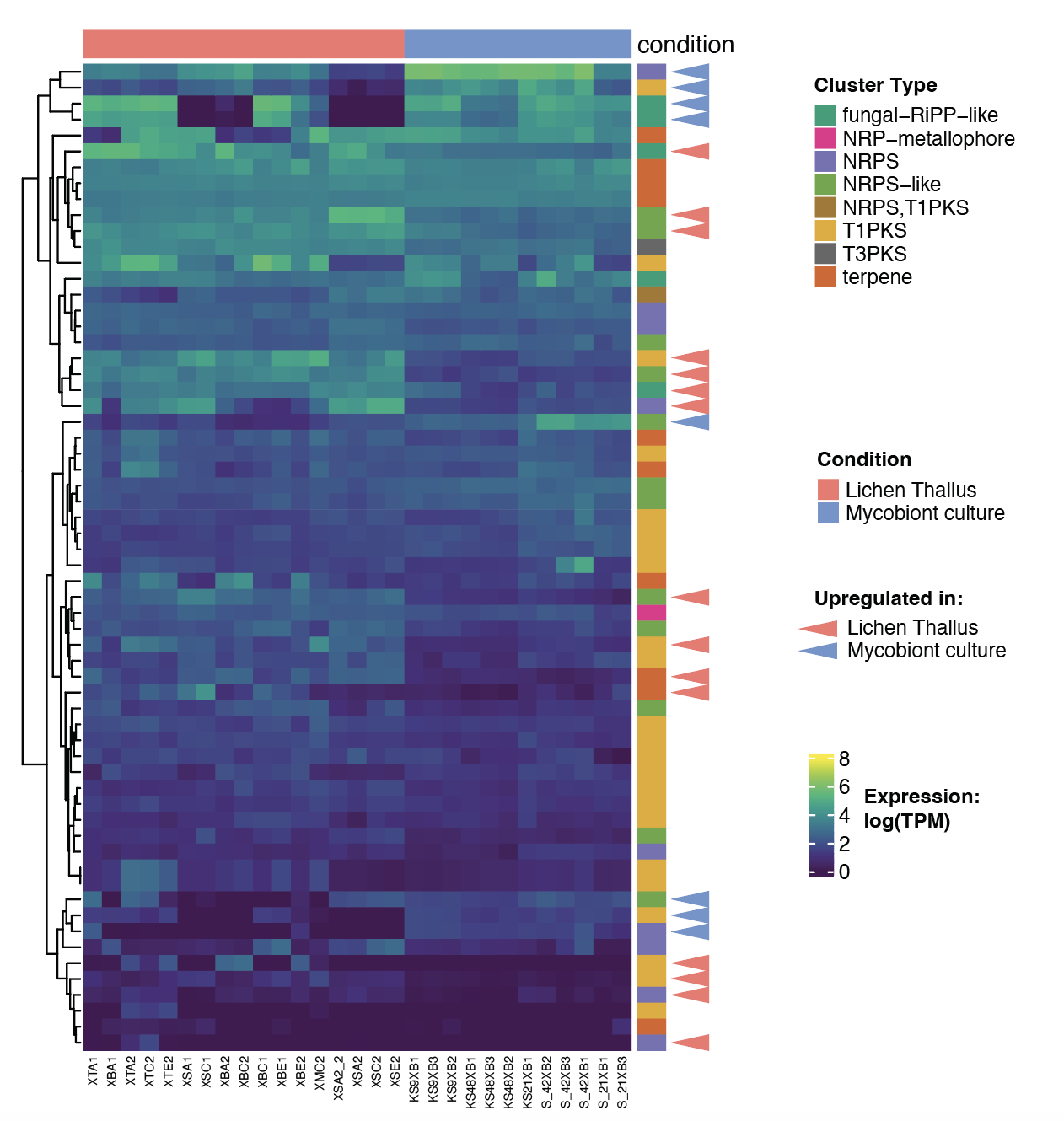

**Figure S3. Differential Gene Expression of core biosynthetic genes in biosynthetic gene clusters of the mycobiont of *X. parietina*.** Heatmap showing expression (as log(TPM), where TPM stands for transcripts per million). The differentially expressed genes are indicated with triangles. The top annotation shows the type of library (lichen or mycobiont culture). The right annotation shows the type of biosynthetic gene cluster.

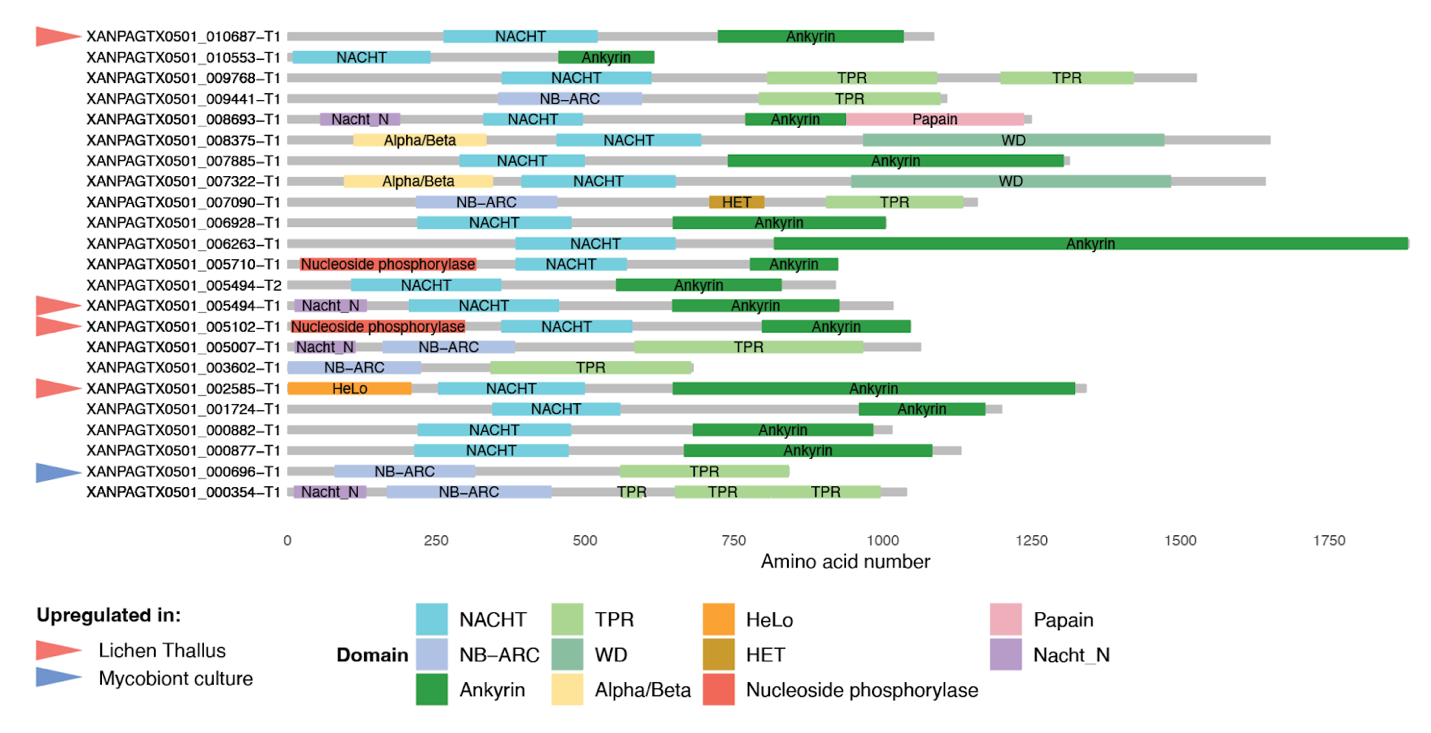

**Figure S4. Putative Nucleotide Oligomerization Domain (NOD)-like receptors (NLRs) in the genome of the mycobiont of *X. parietina.*** In the list of putative NLRs we included protein models which: (a) contained one of the nucleotide-binding domains: NACHT (InterProScan domain: IPR007111), NB-ARC (IPR002182), or AAA (IPR025669); and (b) contained at least one repeat domain: ankyrin (IPR036770, IPR002110, IPR020683), WD40 (IPR001680, IPR036322, IPR015943), or tetratricopeptide repeat (TPR: IPR019734, IPR011990, IPR013026). In the selected proteins, we annotated all other InterProScan domains as well. These included various domains known to act as effector domains in NLRs: Alpha/Beta hydrolases (IPR029058, IPR012908, IPR007751), HeLo domains (IPR038305, IPR029498), Nucleoside phosphorylases (IPR035994, IPR000845), HET domains (IPR010730), and N-terminal domains of NACHT-NTPases (IPR031359, IPR031352). The differentially expressed genes are indicated with triangles.

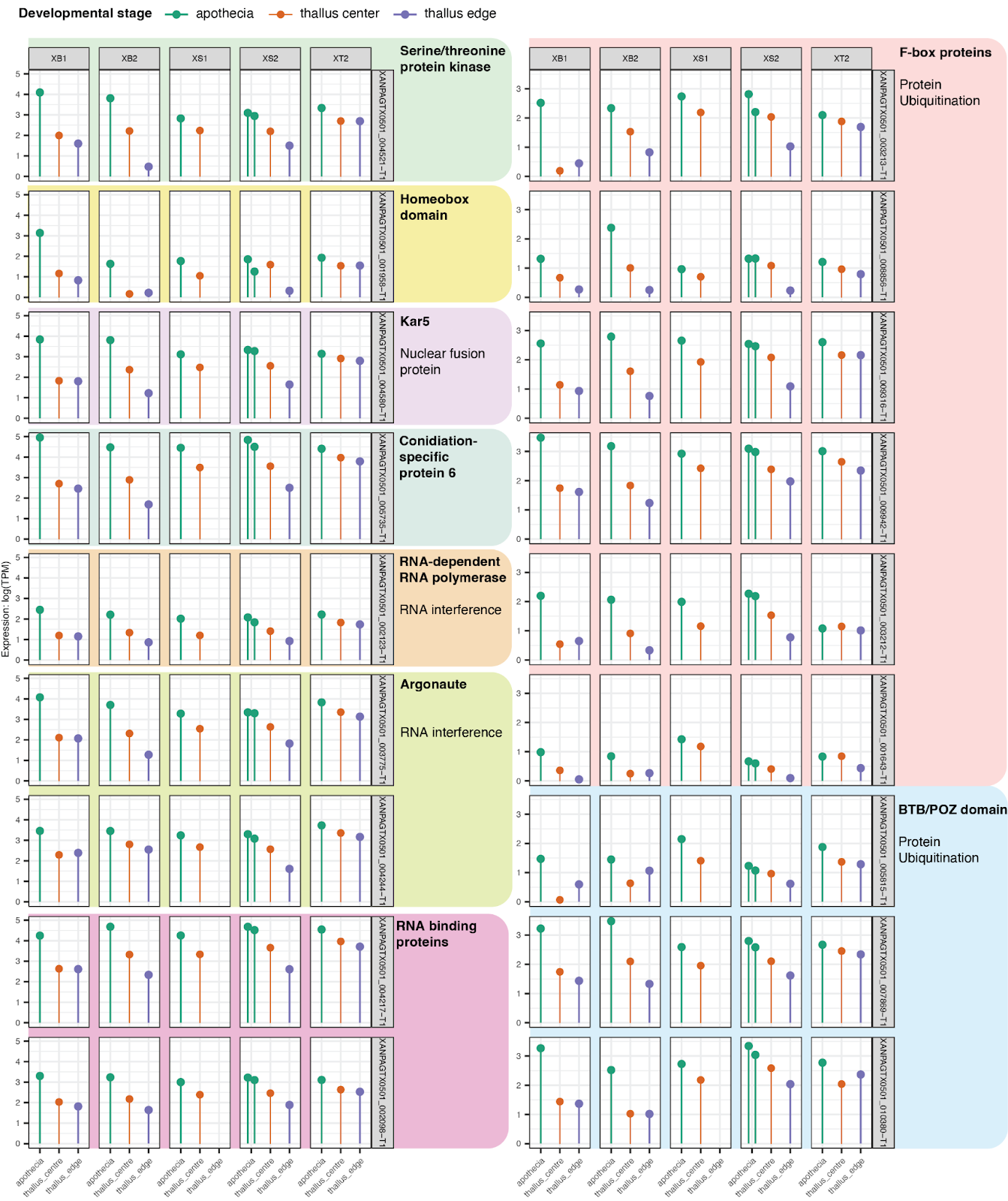

**Figure S5. Expression patterns of several apothecia-upregulated genes in *X. parietina* samples.** The samples are grouped based on the lichen thallus they derived from and colored based on the developmental stage.

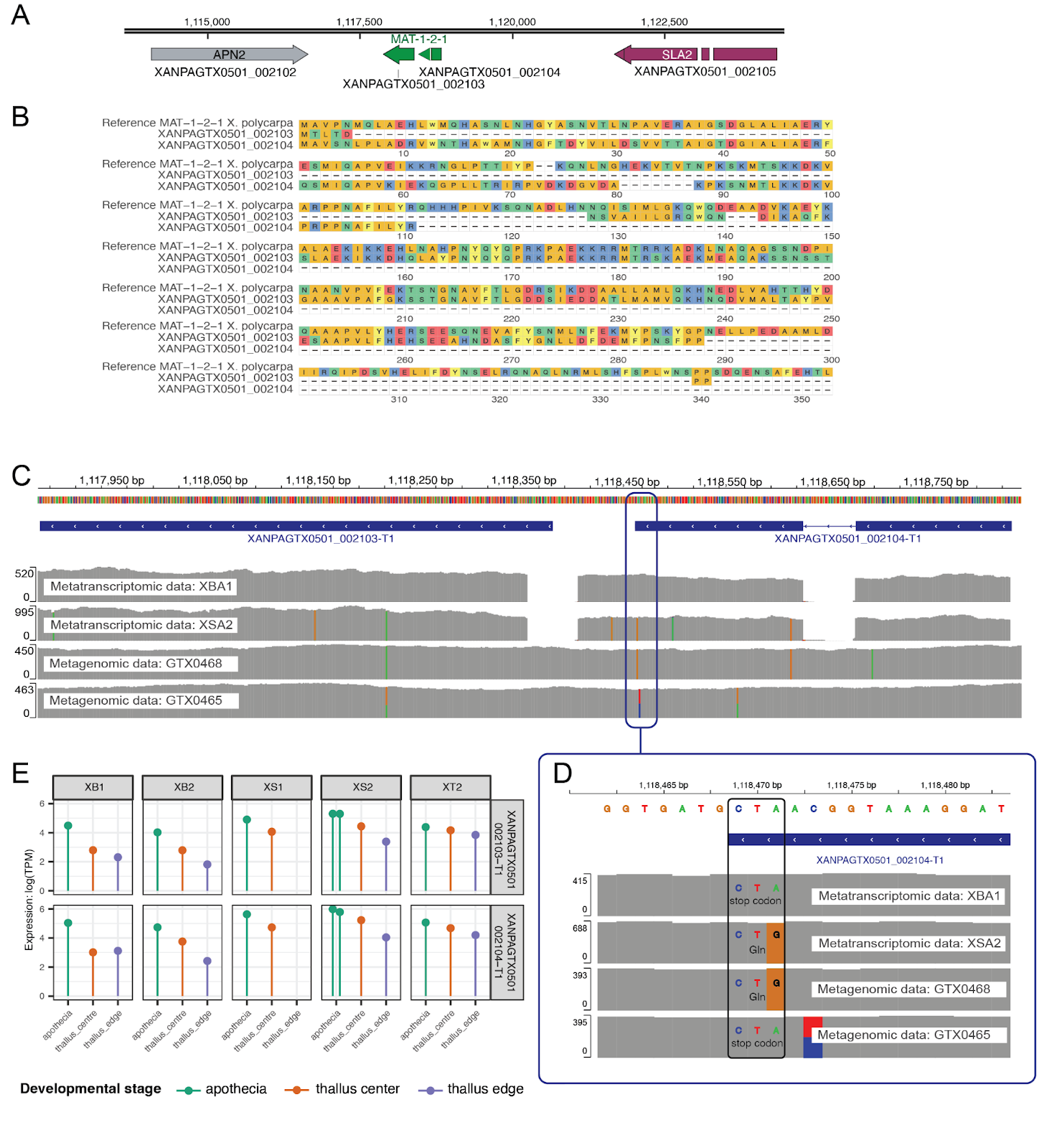
**Figure S6. Mating type locus in *X. parietina*.** To confirm the sex determination systems of the mycobiont, we screened the metagenomic data from eight samples as well as the newly generated genome for presence of both idiomorphs of the mating-type locus. In all generated data, we only detected *MAT-1-2-1*, which is consistent with previous reports and confirms that *X. parietina* has a form of homothallism known as unisexuality  ^134^. Previously Scherrer et al.^135^ reported that *MAT-1-2-1* in *X. parietina*, however, is interrupted by a stop codon. While we confirmed the presence of a stop codon in some of our samples, others lacked it. Moreover, in all samples, *MAT-1-2-1* appears to be expressed and upregulated in apothecia compared to sterile parts of the thallus. This is reminiscent of other unisexual fungi, which often have truncated secondary MAT genes^69^. Future research will determine whether MAT locus is functional in *X. parietina*. A. Structure of the MAT region in *X. parietina*. APN2 and SLA2 flank the MAT locus, which in our annotation was split into two protein models, XANPAGT0501_002103 and XANPAGT0501_002104. B. Alignment of two predicted protein models from *X. parietina*, XANPAGT0501_002103 and XANPAGT0501_002104, and reference sequence of MAT-1-2-1 from *X. polycarpa*. The two protein models align to different parts of the *X. polycarpa* MAT-1-2-1. C. Two metagenomic and two metatranscriptomic libraries mapped onto the MAT-1-2-1 region. The mapping of RNA continues past the stop codon at the end of the gene model XANPAGTX0501_002104. D. The section of C. with the premature stop codon highlighted. While two of the libraries contain the premature stop codon, the two others have a A-> G variant. E. Expression of two transcripts assigned to MAT locus across three different developmental stages of the lichen thallus. The samples are grouped based on the lichen thallus they derived from and colored based on the developmental stage.

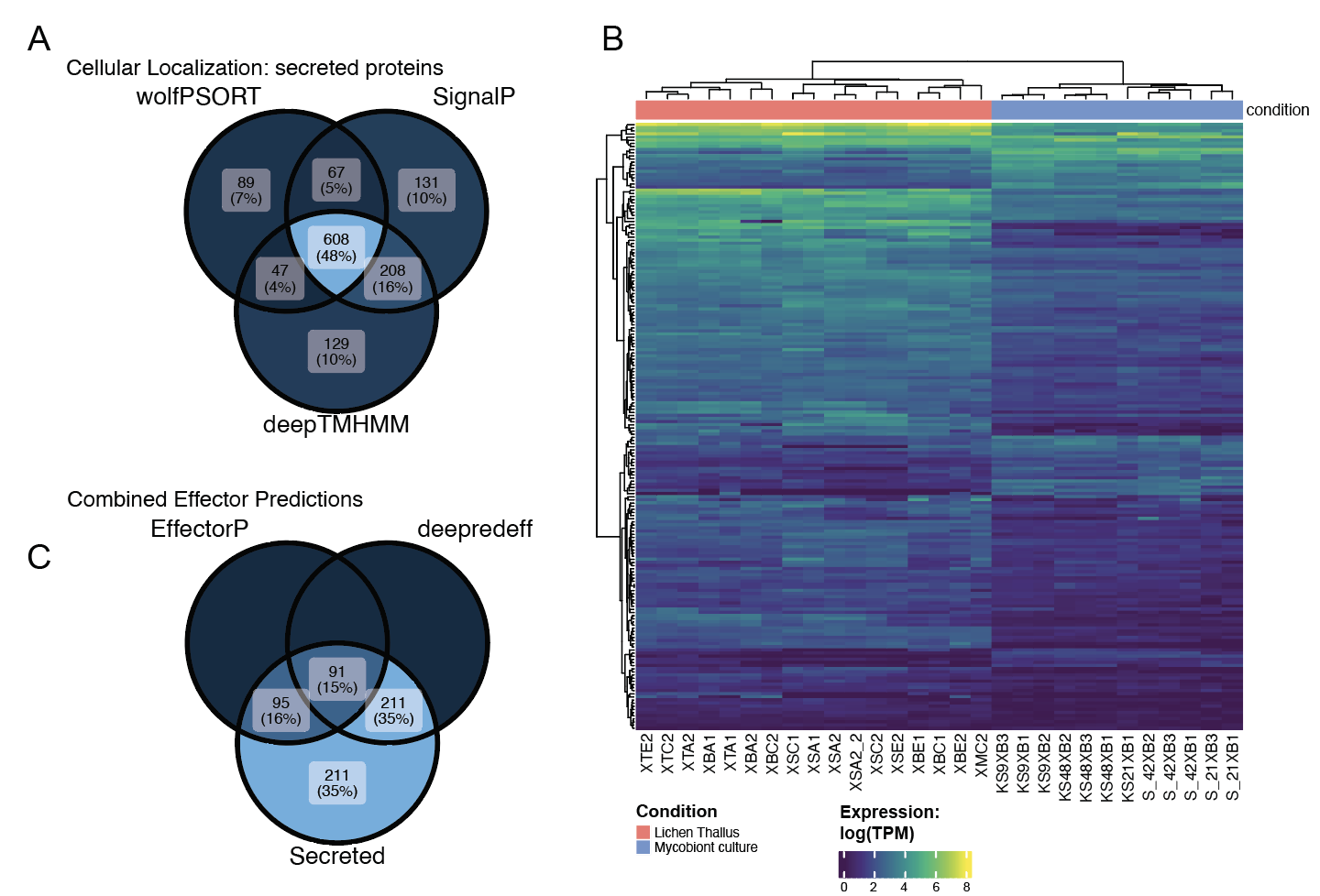

**Figure S7. Prediction and differential expression of the secretome of the mycobiont of *X. parietina.*** A. Venn diagram showing results of three tools used for predicting secreted proteins: SignalP, wolfPSORT, and deepTMHMM. B. Venn diagram showing results of two tools used for effector prediction (EffectorP and deepredeff) juxtaposed with the consensus results of secreted protein prediction (see A). C. Heatmap showing expression levels for the 194 differentially expressed proteins from predicted secretome.
